# Random Forest Regression for Optimizing Variable Planting Rates for Corn and Soybean Using High-Resolution Topographical and Soil Data

**DOI:** 10.1101/2020.02.17.952556

**Authors:** Margaret R. Krause, Savanna Crossman, Todd DuMond, Rodman Lott, Jason Swede, Scott Arliss, Ron Robbins, Daniel Ochs, Michael A. Gore

**Author notes:** Global Wheat Program, International Maize and Wheat Improvement Center (CIMMYT), Ciudad de México, 06600, México (Margaret R. Krause). Corresponding authors: Michael A. Gore, 358 Plant Science Building, Cornell University, Ithaca, NY 14853, USA,; Margaret R. Krause, Centro Internacional de Mejoramiento de Maíz y Trigo, Apdo. Postal 6-641 06600 Ciudad de México., MÉXICO.

## Abstract

In recent years, planting machinery that enables precise control of the planting rates has become available for corn (*Zea mays* L.) and soybean (*Glycine max* L.). With increasingly available topographical and soil information, there is a growing interest in developing variable rate planting strategies to exploit variation in the agri-landscape in order to maximize production. A random forest regression-based approach was developed to model the interactions between planting rate, topography, and soil characteristics and their effects on yield based on on-farm variable rate planting trials for corn and soybean conducted at 27 sites in New York between 2014 and 2018 (57 site-years) in collaboration with the New York Corn and Soybean Growers Association. Planting rate ranked highly in terms of random forest regression variable importance while explaining relatively minimal yield variation in the linear context, indicating that yield response to planting rate likely depends on complex interactions with agri-landscape features. Models were moderately predictive of yield within site-years and across years at a particular site, while the ability to predict yield across sites was low. Relatedly, variable importance measures for the topographical and soil features varied considerably across sites. Together, these results suggest that local testing may provide the most accurate optimized planting rate designs due to the unique set of conditions at each site. The proposed method was extended to identify the optimal variable rate planting design for maximizing yield at each site given the topographical and soil data, and empirical validation of the resulting designs is currently underway.

## INTRODUCTION

In agricultural production systems, site-specific management involves the development of crop management strategies at a finer spatial scale than that of the whole field area (Plant, 2001). Compared to early farming, which was performed by hand and thereby facilitated site-specific management, the advent of mechanization in farming ushered in an era of uniform application of inputs on large areas of cropland. Studies have shown that uniform applications can result in suboptimal input use efficiency in parts of the field that may require greater or fewer inputs than the applied fixed rated (Mulla & Schepers, 1997; Moore & Tyndale-Biscoe, 1999), though increases in farm productivity generally outweighed the associated economic losses. However, the growing costs of agricultural inputs (USDA NASS, 2017) and negative environmental impacts of intensive production practices (Tilman et al., 2002) threaten the long-term economic and environmental sustainability of uniform crop management.

Variable rate application systems enable growers to apply inputs at a range of user-defined levels within a field area. Variable rate as it applies to planting is not a new concept. The first variable rate planting systems emerged in the United States during the 1970s but did not become popular due to their initial lack of automation, which necessitated that growers trigger planting rate transitions manually (Lowenberg-DeBoer, 1999). In the late 1990s, the integration of GPS and automated variable rate systems for commercial planting equipment renewed interest in variable rate planting technologies.

However, these improvements did not immediately drive a widespread adoption of variable rate planting, as users remained uncertain of how to develop variable rate planting designs to maximize returns given the underlying field conditions. Some early as well as more recent studies have based the development of variable rate planting designs on yield potential, delineated by previous years’ yield performance and/or the grower’s knowledge of the field (Barnhisel et al. 1996; Hörbe et al., 2013; Corassa et al., 2018). However, Bullock et al. (1998) postulated that knowledge of yield potential would be insufficient to make variable rate technologies economically beneficial to farmers. They identified as a major barrier the community’s limited understanding of how planting rate and various field characteristics such as topography and soil which can be difficult to measure may interact to influence yield. More than 20 years later, use of variable rate planting technology remains limited, despite the increasing availability of high-resolution topographical and soil data and the abundance of commercial variable rate products on the market. In a 2016 survey of corn and soybean growers in New York State, only 10.5 percent of respondents had adopted variable rate planting technology and 20.0 percent cited skepticism towards the technology’s potential economic benefit as a key driver of its slow adoption (van Es et al., 2016).

Currently, there is little consensus among the community regarding how to create variable rate planting designs with respect to yield potential, field characteristics, and/or other observations. While many research efforts have attempted to determine the relationships between topographic features, soil characteristics, and yield potential for crops sown under fixed planting rates (Miller et al., 1988; Katerji et al., 1995; Changere & Lal, 1997; Kravchenko & Bullock, 2000; Frogbrook et al., 2002; Cox et al., 2003), few studies have assessed how planting rate might influence and interact with those relationships, and among those, the results have been mixed: some have observed potential for the profitability for variable rate planting technology (Shanahan et al., 2004), while others found inconsistencies among planting rate optimizations and the interactions between planting rate and topographical/soil features across sites and years (Smidt et al. 2016; Licht et al. 2017).

In a literature review of variable rate technologies, Bullock and Lowenberg-DeBoer (2007) made the observation that the majority of studies pertaining to variable rate planting and chemical applications have reported results for a small number of specific site-years. The authors, among others (Lambert et al., 2006; Liu et al. 2006; Bullock & Lowenberg-DeBoer, 2007; Ruffo et al. 2006), emphasized the need for longer-term, multi-location experimental data in order to expand the inference space and sample the population of possible environmental conditions at a high level in order to evaluate the value and feasibility of variable rate planting.

To conduct experiments on such a scale, coordination between growers and researchers are needed in addition to a long-term commitment to on-farm testing. The New York Corn and Soybean Growers Association (NYCSGA) is a statewide non-profit organization that represents the interests of New York corn and soybean producers and sponsors relevant research on production, utilization, and marketing. In 2013, the NYCSGA developed a research initiative to optimize variable rate planting technology for corn and soybean growers in New York State. NYCSGA members throughout central, western, and northern New York conducted on-farm variable rate field trials between 2014 and 2018 with the objective of developing a strategy for variable rate planting to exploit the extensive native variation of New York’s agricultural landscape. To sample this variation, spatial data related to topographical features, soil type, and soil nutrients were collected at a high resolution.

This collaborative project driven by growers has created a wealth of information relating yield to planting rate and its interactions with a wide range of environmental factors. However, the magnitude and complexity of these data present a number of statistical challenges when conducting analysis. Many oft-recorded variables in environmental or agricultural studies exhibit significant correlations among one another. For example, topographical attributes such as elevation and slope can influence the movement of water through and over the landscape, thereby affecting soil development (Moore et al. 1993). Relationships between topographical features and soil characteristics such as sand, silt, organic matter, pH, cation exchange capacity, and extractable phosphorus have been reported (Brubaker et al., 1993; Moore et al., 1993; Tan et al., 2004). Likewise, strong relationships between soil variables themselves are not uncommon. In a linear-regression analysis of the effects of soil characteristics on corn yield response to variable planting rates, Licht et al. (2017) removed variables describing silt and available water-holding capacity due to their collinearity with sand, clay, and soil organic matter. In addition to correlations between variables, nonlinear relationships can often occur in agricultural systems (reviewed in Archontoulis & Miguez, 2014). For example, increasing temperatures are known to benefit corn yields but become harmful once they reach 30°C and above (Schlenker & Roberts, 2006). Despite the limited ability of linear regression to accurately model correlated characters and nonlinear relationships, many published studies evaluating the potential of variable rate planting technology have used it to assess the relationships between planting rate, yield, and environmental features (Shanahan et al., 2004; Hörbe et al., 2013; Licht et al., 2017).

Novel statistical approaches to analyze increasingly large and complex agronomic datasets are needed to fully exploit the information they contain. Random forest regression is an ensemble learning, nonparametric method that constructs multiple decision trees using random subsets of the observations and predictor variables (Breiman, 2001). Random forest regression models have become increasingly popular in a range of scientific fields due to their ability to accommodate highly correlated variables, complex nonlinear interactions, and predictors with varying types and ranges. Random forest regression has been used effectively for environmental and agricultural applications such as predicting soil texture from terrain variables (Ließ et al., 2012) and estimating vegetative biomass from remotely sensed data (Mutanga et al., 2012). This method may also be useful for capturing the complex relationships between planting rate, yield, and field characteristics with the goal of developing optimized variable rate planting designs.

The work presented here aims to 1) investigate the effects of planting rate, topographical features, soil characteristics, and their interactions on corn and soybean yield and 2) propose a random forest regression-based method for leveraging this information to build variable rate planting designs.

## MATERIALS AND METHODS

### Site-Year Summary

On-farm corn and soybean variable rate field trials were conducted in 27 unique sites throughout central, western, and northern New York (Fig. 1) between the years 2014 and 2018 for a total of 57 site-years (32 corn, 25 soybean). Field area, row spacing, experimental design, planting rates tested, and hybrids/varieties varied among site-years (Table 1). Field areas ranged from 10.8 to 59.9 ha with an average of 28.0 ha. The row spacings used were 50.8, 76.2, 101.6, and 127.0 cm. For corn, 16 hybrids were used throughout the course of the experiment, though hybrids P0533AM1 and P0216/P0216AM were used most predominantly, appearing in 15 and 8 of the 32 site-years, respectively. For soybean, 21 varieties were used. Statewide weather data were obtained by the National Climatic Data Center (National Climatic Data Center, 2019) (Fig. 2).

**FIGURE 1.**
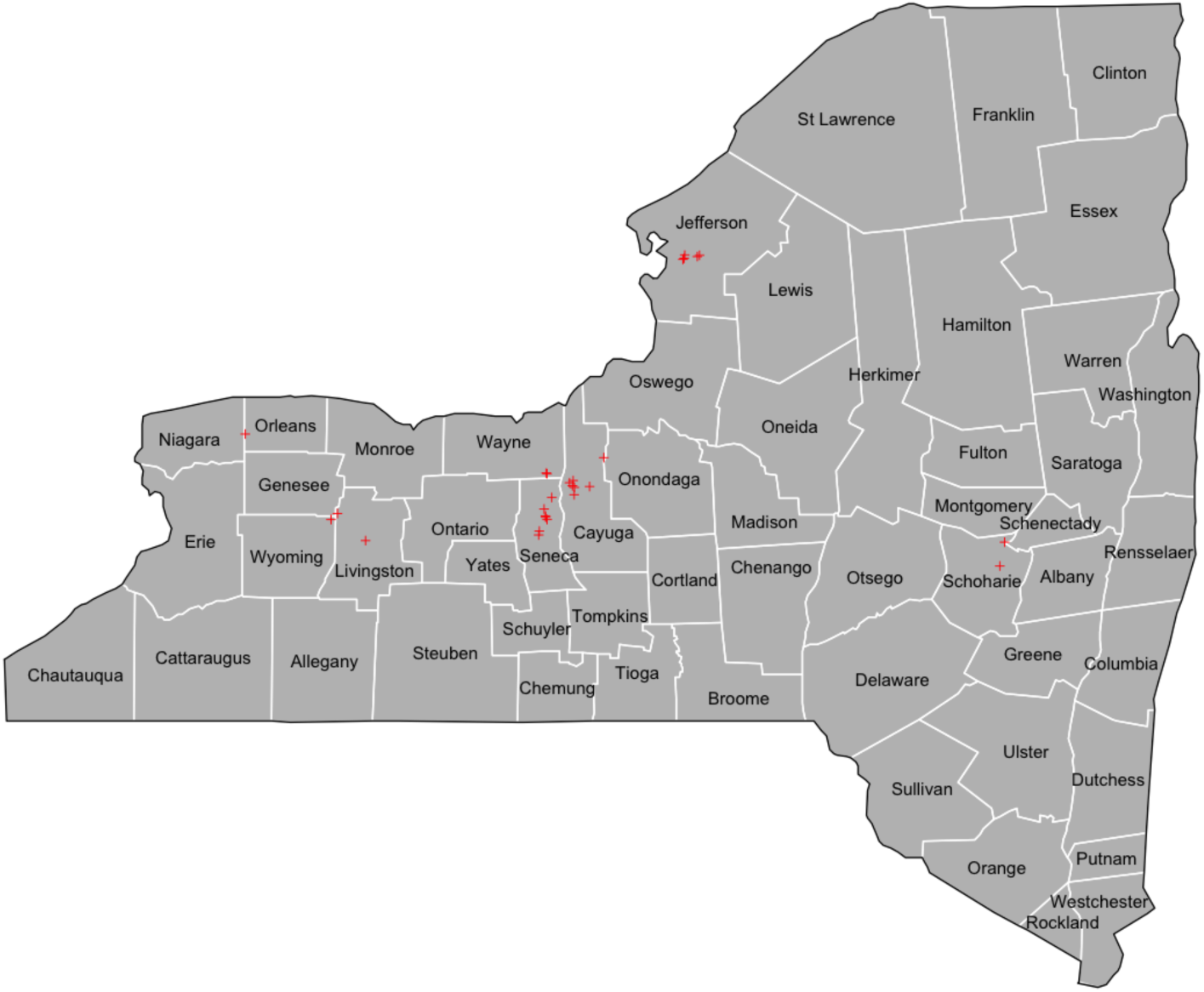
County-level map of the sites in which on-farm variable rate trials were conducted between 2014 and 2018 in New York

**FIGURE 2.**
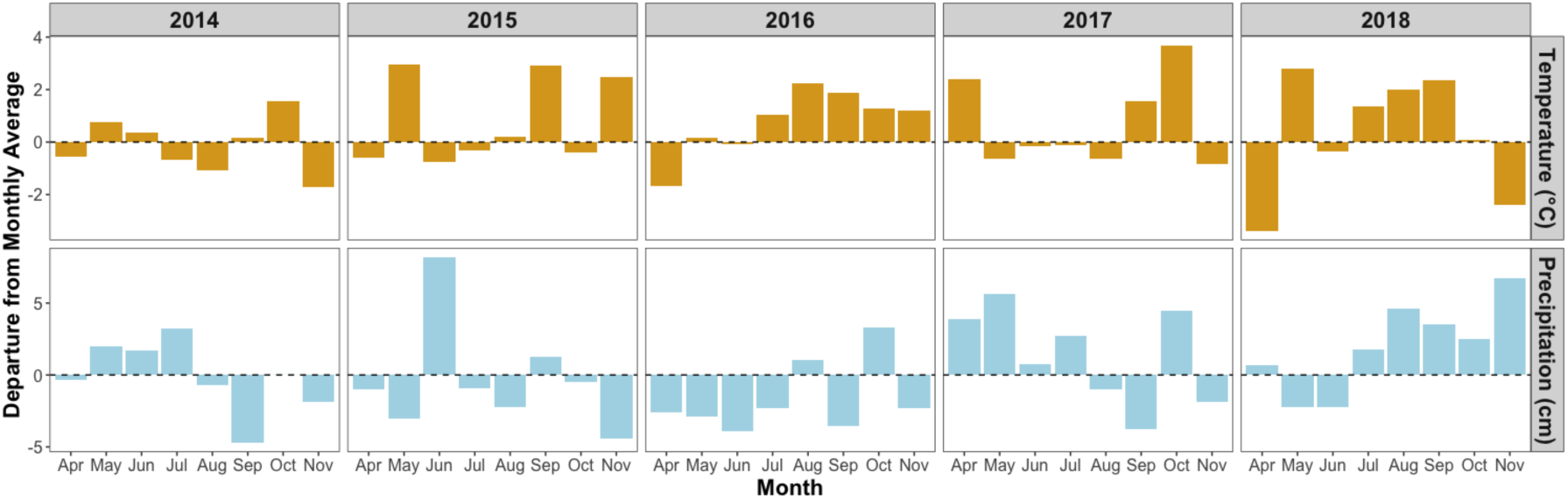
Statewide departures from 1981-2010 monthly temperature and precipitation averages during the 2014-2018 growing seasons

**TABLE 1.** Site-year-specific information for each on-farm field trial

| Field | Year | Crop | Area (ha) | Row Spacing (cm) | Hybrid/Variety | Planting Rate Design | Hybrid/Variety Design |
| --- | --- | --- | --- | --- | --- | --- | --- |
| An6 | 2014 | Corn | 22.81 | 50.8 | P9690AM<br>P9675AMXT | Randomized Blocks | Alternating Hybrids/Varieties |
| An8 | 2014 | Corn | 29.99 | 50.8 | P0216<br>P0533AM1 | Randomized Blocks | Alternating Hybrids/Varieties |
| An8 | 2015 | Corn | 29.99 | 50.8 | P0216<br>P0533AM1 | Randomized Blocks | Alternating Hybrids/Varieties |
| Ba1 | 2015 | Corn | 21.23 | 76.2 | P0216<br>P0533AM1 | Randomized Blocks | Alternating Hybrids/Varieties |
| Be2 | 2014 | Corn | 51.15 | 76.2 | P0216<br>P0533AM1 | Randomized Blocks | Alternating Hybrids/Varieties |
| Bi | 2015 | Corn | 14.92 | 101.6 | P0533AM1 | Randomized Blocks | Single Hybrid/Variety |
| Bi | 2016 | Corn | 14.92 | 101.6 | P0157AMX | Randomized Blocks | Single Hybrid/Variety |
| Bi | 2016 | Corn | 14.92 | 101.6 | 197-68STXRIB | Randomized Blocks | Single Hybrid/Variety |
| Cr | 2014 | Corn | 29.55 | 101.6 | DKC46-20RIB<br>P0094AMX | Randomized Blocks | Alternating Hybrids/Varieties |
| Cr | 2015 | Corn | 29.55 | 101.6 | P0216AM | Randomized Blocks | Single Hybrid/Variety |
| Dm2 | 2014 | Corn | 23.47 | 50.8 | P0216AM<br>P0533AM1 | Randomized Blocks | Alternating Hybrids/Varieties |
| Du1 | 2014 | Corn | 24.34 | 76.2 | P0216AM<br>P0533AM1 | Randomized Blocks | Alternating Hybrids/Varieties |
| Du3 | 2016 | Corn | 39.63 | 76.2 | P0216AM<br>P0533XR | Randomized Blocks | Alternating Hybrids/Varieties |
| He15 | 2015 | Corn | 23.75 | 50.8 | P0157AMX<br>DK52-85 | Randomized Blocks | Alternating Hybrids/Varieties |
| Hk | 2015 | Corn | 35.07 | 50.8 | P0533<br>P0604 | Randomized Blocks | Alternating Hybrids/Varieties |
| Ke | 2015 | Corn | 31.31 | 76.2 | P0216AM<br>P0533AM1 | Randomized Blocks | Alternating Hybrids/Varieties |
| Mc2 | 2015 | Corn | 15.83 | 50.8 | P0604<br>P0533AM1 | Randomized Blocks | Alternating Hybrids/Varieties |
| Mc3 | 2015 | Corn | 17.78 | 50.8 | P0506AM<br>P0533AM1 | Randomized Blocks | Alternating Hybrids/Varieties |
| Ov29 | 2014 | Corn | 23.97 | 101.6 | P0094AMX | Randomized Blocks | Single Hybrid/Variety |
| Ov29 | 2015 | Corn | 23.97 | 101.6 | P0094AMX | Randomized Blocks | Single Hybrid/Variety |
| Ov30 | 2014 | Corn | 10.80 | 101.6 | P0094AMX | Randomized Blocks | Single Hybrid/Variety |
| Ov30 | 2015 | Corn | 10.80 | 101.6 | P0604AM | Randomized Blocks | Single Hybrid/Variety |
| Ri23 | 2015 | Corn | 40.71 | 76.2 | P0216AM<br>P0533AM1 | Randomized Blocks | Alternating Hybrids/Varieties |
| Ri27 | 2015 | Corn | 12.74 | 50.8 | P0216AM<br>P0533AM1 | Randomized Blocks | Alternating Hybrids/Varieties |
| St | 2015 | Corn | 47.71 | 76.2 | P0216AM<br>P0533AM1 | Randomized Blocks | Alternating Hybrids/Varieties |
| Sc | 2015 | Corn | 23.71 | 101.6 | P0157AMX | Randomized Blocks | Single Hybrid/Variety |
| Sc | 2016 | Corn | 23.71 | 101.6 | P0533AM1 | Randomized Blocks | Single Hybrid/Variety |
| Sc | 2017 | Corn | 23.71 | 101.6 | P0157AMX | Randomized Blocks | Single Hybrid/Variety |
| Sw37 | 2015 | Corn | 23.72 | 76.2 | 49-72 | Randomized Blocks | Single Hybrid/Variety |
| Sw52 | 2016 | Corn | 19.07 | 76.2 | 49-72 | Randomized Blocks | Single Hybrid/Variety |
| Sw52 | 2017 | Corn | 19.07 | 76.2 | 49-72 | Randomized Blocks | Single Hybrid/Variety |
| Vc1 | 2016 | Corn | 59.93 | 50.8 | P0506AM<br>P0533AM1<br>P0157AMX | Randomized Blocks | Alternating Hybrids/Varieties |
| An6 | 2015 | Soybean | 22.81 | 101.6 | 2108R | Randomized Blocks | Single Hybrid/Variety |
| Ba1 | 2016 | Soybean | 21.23 | 101.6 | AG17X6 | Alternating Strips | Single Hybrid/Variety |
| Do | 2015 | Soybean | 56.64 | 101.6 | AG3034 | Randomized Blocks | Single Hybrid/Variety |
| Do | 2016 | Soybean | 56.64 | 101.6 | AG3334 | Randomized Blocks | Single Hybrid/Variety |
| Du1 | 2015 | Soybean | 24.34 | 101.6 | P24T19R | Alternating Strips | Single Hybrid/Variety |
| Du3 | 2015 | Soybean | 39.63 | 101.6 | P24T19R<br>92Y91 | Alternating Strips | Sections |
| He15 | 2014 | Soybean | 23.75 | 101.6 | P92Y12<br>P92Y51 | Randomized Blocks | Alternating Hybrids/Varieties |
| He15 | 2016 | Soybean | 23.75 | 101.6 | P24T05 | Randomized Blocks | Single Hybrid/Variety |
| Hk | 2014 | Soybean | 35.07 | 127 | P92Y51 | Randomized Blocks | Single Hybrid/Variety |
| Ke | 2016 | Soybean | 31.31 | 101.6 | P19T01R | Alternating Strips | Single Hybrid/Variety |
| Lb | 2016 | Soybean | 28.53 | 50.8 | P19T/P01T<br>P19T04R 15 | Randomized Blocks | Sections |
| Mc1 | 2014 | Soybean | 25.17 | 101.6 | AG3030 | Randomized Blocks | Single Hybrid/Variety |
| Mc1 | 2018 | Soybean | 25.17 | 101.6 | AG3334<br>SG1776 | Randomized Blocks | Sections |
| Mc2 | 2014 | Soybean | 15.83 | 101.6 | P93Y22 | Randomized Blocks | Single Hybrid/Variety |
| Mc2 | 2016 | Soybean | 15.83 | 101.6 | AG2035 | Randomized Blocks | Single Hybrid/Variety |
| Mc3 | 2014 | Soybean | 17.78 | 101.6 | AG3030 | Randomized Blocks | Single Hybrid/Variety |
| Ov29 | 2018 | Soybean | 23.97 | 76.2 | SG1863XT<br>P16A35X | Randomized Blocks | Sections |
| Ov30 | 2018 | Soybean | 10.80 | 76.2 | SG1863XT<br>P16A35X | Randomized Blocks | Sections |
| St | 2014 | Soybean | 47.71 | 101.6 | P92Y51 | Alternating Strips | Single Hybrid/Variety |
| Sw37 | 2014 | Soybean | 23.72 | 76.2 | 2031 | Randomized Blocks | Single Hybrid/Variety |
| Sw37 | 2016 | Soybean | 23.72 | 76.2 | AG2031 | Randomized Blocks | Single Hybrid/Variety |
| Sw52 | 2015 | Soybean | 19.07 | 76.2 | AG2431 | Randomized Blocks | Single Hybrid/Variety |
| Tp80 | 2016 | Soybean | 31.27 | 76.2 | Unknown | Randomized Blocks | Single Hybrid/Variety |
| Vc1 | 2015 | Soybean | 59.93 | 101.6 | P22T41<br>P92Y51 | Randomized Blocks | Sections |
| Vc1 | 2017 | Soybean | 59.93 | 101.6 | P22T41<br>AG2035 | Randomized Blocks | Sections |

### On-Farm Experimental Designs

All on-farm variable rate trials for corn were sown with a randomized design consisting of 0.81 ha (2 ac) blocks sown to one of four target planting rates (Fig. 3). For corn, the following four target planting rates were tested: 66,718 (27,000), 79,074 (32,000), 91,429 (37,000), and 103,784 (42,000) seeds ha^-1^ (ac^-1^). Out of the 32 site-years for corn, 17 were sown with two hybrids and 1 was sown with three hybrids. All on-farm trials containing more than one hybrid were sown with a split planter, creating alternating strips of each hybrid within the field.

**FIGURE 3.**
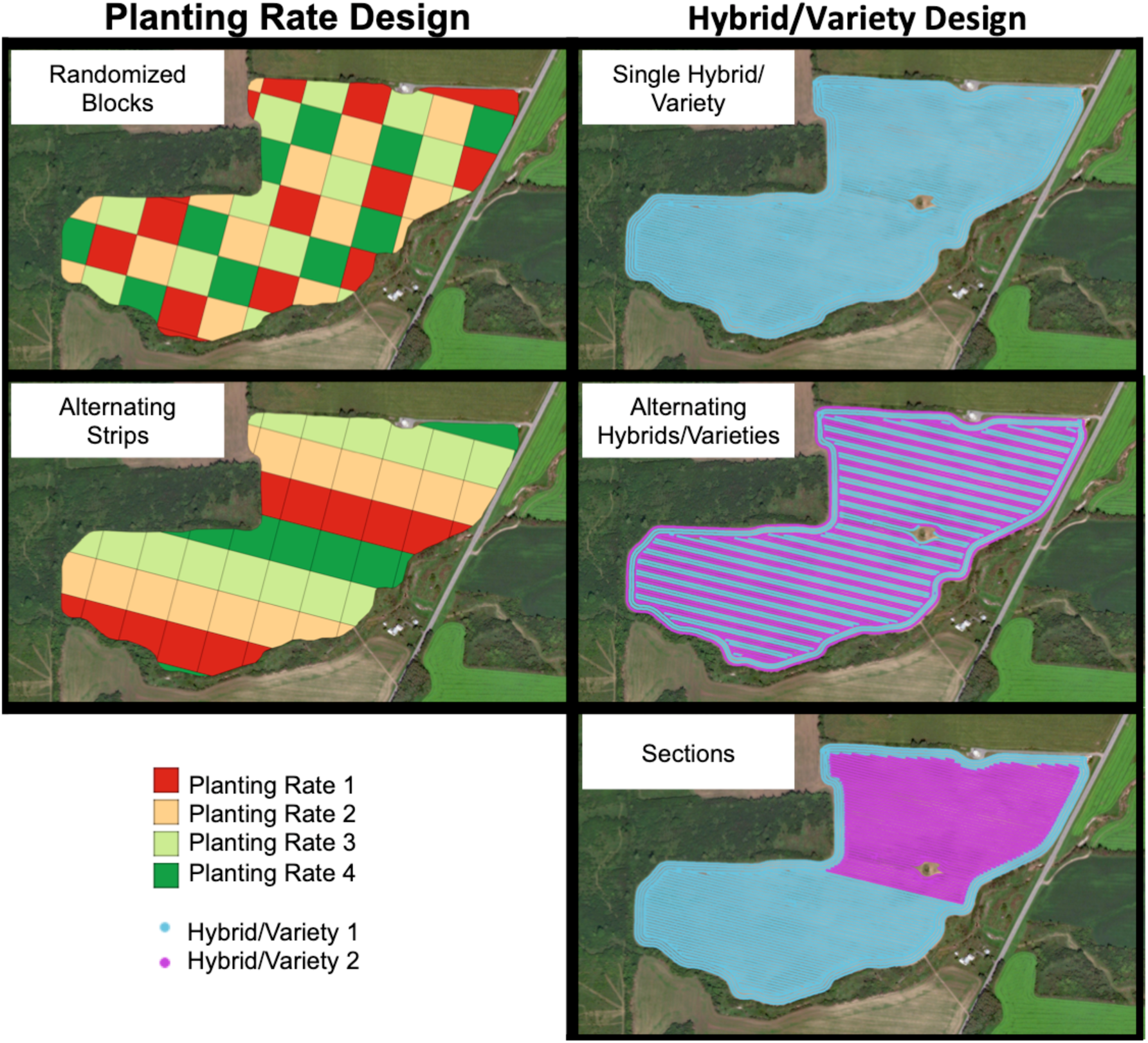
Field designs for planting rate and hybrid/variety

For soybean, the same 0.81 ha target planting rate block system was used with the following target rates: 197,684 (80,000), 296,526 (120,000), 395,368 (160,000), and 494,210 (200,000) seeds ha^-1^ (ac^-1^). Rates were assigned to blocks at random for 20 of the 25 site-years. Because variable rate soybean planters were not available for the remaining 5 site-years, target planting rates were assigned to the rows or columns of the blocks, enabling adjustments to be made to the planting rate between passes of the planter. A total of 7 of the 25 site-years were sown with two soybean varieties. A split soybean planter was only available for site-year He15_2014, which was sown with alternating varieties. The remaining 6 site-years were sown such that each variety occupied a section of the field (Fig. 3)

### Experimental Unit Grid

The 0.81 ha target planting rate block system was developed to minimize error associated with transitioning between planting rates. There are, however, limitations to treating the 0.81 ha target planting rate blocks as the experimental units. Blocks that fell along the edges of the field were often irregularly shaped (as seen in Fig. 3). As a result, the number of full-sized 0.81 ha blocks in smaller fields was low and insufficient for analysis. Furthermore, due to the high variability of the topographical and soil features observed in some fields, some of the 0.81 ha blocks contained a wide range of variability. Lastly, because split planters were used to sow multiple hybrids/varieties in some site-years, individual blocks contained more than one hybrid/variety. Therefore, by using the blocks as the experimental units, it would not be possible to evaluate the interactions of hybrid/variety with planting rate, soil features, and topographical characteristics and their effects on yield.

To address these issues, a finer resolution grid-based system was developed to serve as the experimental units. Square grids were created in QGIS (QGIS Development Team, 2019) with the length and width of the grid cells set equal to the width of the planter, unless a split planter was used to sow multiple hybrids/varieties (Fig. 4). In that case, the length and width of the grid cells were set to half the width of the planter. Each grid cell was assigned a unique identifier (ID). All data types were aggregated to the experimental unit grid to resolve the data into a tabular format for analysis.

**FIGURE 4.**
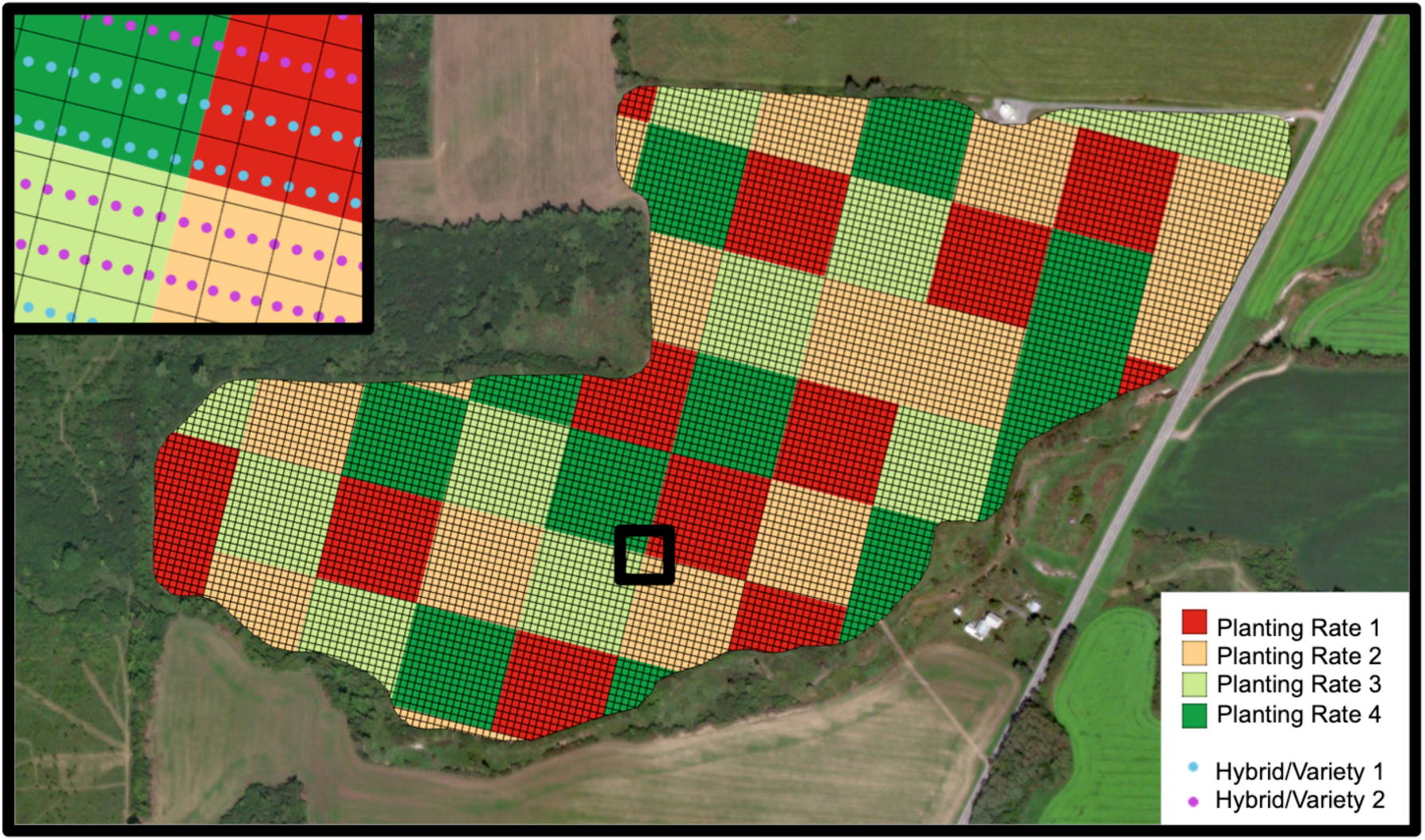
Grid of experimental units with respect to the randomized block planting rate design and the alternating hybrids/varieties design The black box within the field denotes the area represented in the inlay. The length and width of the grid cells in this example are equal to half the width of the split planter such that data from only one hybrid/variety are present within each cell.

### Data Types and Quality Control

For each site-year, five spatial data layers were used to assess the relationship between yield, planting rate, hybrid/variety, topographical characteristics, and soil characteristics. The layers were as follows: 1) harvest, 2) target/as-applied planting rate 3) topography, 4) grid soil sampling, and 5) Natural Resources Conservation Service (NRCS) Soil Survey (Fig. 5).

**FIGURE 5.**
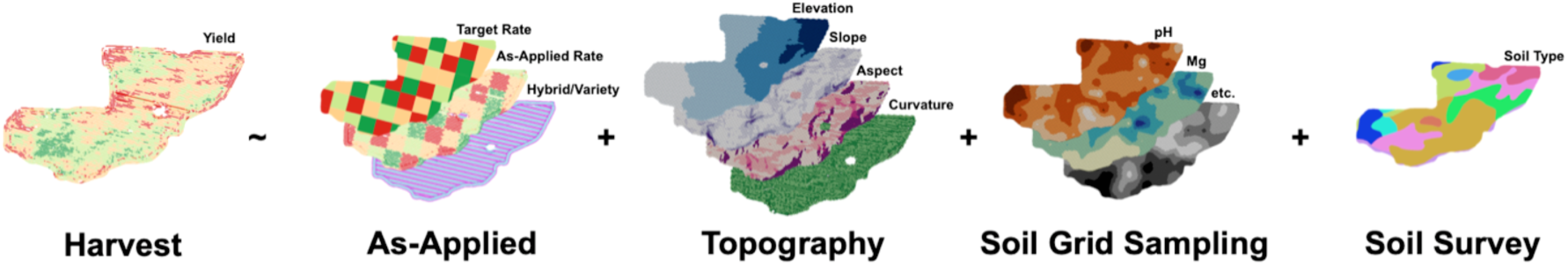
The five spatial data types used to model the relationship between yield, planting rate, hybrid/variety, topographical features, and soil characteristics

The harvest point layer contained the levels of dry volume yield harvested throughout the field. In the form of polygons, the target planting rate layer contained the randomized 0.81 ha block design with which the trials were sown, while the as-applied planting rate point layer recorded the precise rates applied by the planter as well as where each hybrid/variety was sown. These three layers were used in conjunction with one another to perform various quality control measures to the as-applied and yield data (Fig. S1A-F). Briefly, the headlands were removed from both the as-applied planting and harvest layers in QGIS. The remaining data points were then assigned ID numbers corresponding to the experimental unit grid cell in which they appeared. The as-applied planting and harvest layers were then analyzed in R (R Core Team, 2019) to remove outlying grid cells based on the mean, variance, and number of data points within each cell. Grid cells were also removed from the analysis if they contained data corresponding to more than one target planting rate or hybrid/variety. Finally, grid cells in which the as-applied planting rate deviated from the target planting rate beyond a threshold of ±1011.7 seeds ha^-1^ for corn and ±8093.7 seeds ha^-1^ for soybean were also removed.

Topographical variables for elevation, slope, aspect, and curvature were derived from elevation measurements recorded by the GPS systems aboard the planting machinery (Fig. S1G). Using the as-applied planting rate layer, elevation means were calculated in QGIS using all of the data points within each experimental grid cell. Slope, aspect, and curvature were then calculated in R using 3×3 neighborhoods of cells following Burrough and McDonell (1998) and Zevenbergen and Thorne (1987).

Integrated Ag Services (Mildford Center, OH) conducted grid soil sampling along a 0.20-ha grid pattern, and Spectrum Analytic (Washington Court House, OH) provided the topsoil analysis. Point data on up to 12 soil features were provided for each site: aluminum (Al), phosphorus (P), potassium (K), potassium saturation (KSt), calcium (Ca), calcium saturation (CaSt), magnesium (Mg), magnesium saturation (MgSt), pH, buffer pH (BpH), cation exchange capacity (CEC), and soil organic matter (OM). K, Mg, and CEC were recorded for all 27 sites (57 site-years). The remaining variables were unavailable for ≤ 4 sites (≤ 12 site-years) with the exception of Al, which was available for only 17 of the 27 sites (37 site-years). Using the “gstat” package in R, the soil sample measurements were block kriged to the experimental unit grid (Fig. S1G). If a semivariogram for a particular variable could not be fit due to low spatial autocorrelation, the values for that variable were set to missing.

The NRCS Soil Survey layer in the form of polygons was obtained for each sites using the SSURGO database (Soil Survey Staff, Natural Resources Conservation Service, USDA, 2015). The soil type attribute was extracted for each experimental grid cell using QGIS. If an experimental unit grid cell spanned multiple NRCS Soil Survey polygons with differing reported soil types, the soil type variable for that cell was set to missing (Fig. S1I)

Final datasets containing variables describing the yield, target and as-applied planting rates, topography, grid soil sampling, and NRCS Soil Survey were merged into a combined data frame in R according to the experimental unit grid cell IDs. Excluding yield as the response variable, the dimensions of the final datasets for each site-year used for model fitting were *n* × *p* where *n* is the number of experimental unit grid cells and *p* is the number of predictors.

### Variable Correlation, ANOVA, and Linear Regression

Pearson’s correlations were calculated between all variables. Treatment effects for planting rate and hybrid/variety were estimated using ANOVA. Yield was regressed on each topographical and soil variable to estimate the percent of yield variation explained (*R^2^*). Yield was also regressed on planting rate, hybrid/variety, and their interaction to assess their collective influence on yield in a linear context. All correlation, ANOVA, and linear regression analyses were carried out in R.

### Random Forest Regression

Random forest regression models were fit to assess prediction accuracy, calculate variable importance, and develop optimized planting rate designs. Forests were grown using the “cforest” function of the “party” package for R (Hothorn et al. 2006; Strobl et al. 2007; Strobl et al. 2008). Through simulation, Strobl et al. (2007) demonstrated that bias could occur in the variable importance measures when predictors vary in their scale of measurement or, for categorical variables, in their number of levels. To avoid bias, Strobl et al. (2007) proposed “cforest” as an alternative implementation of the commonly used regression functions in the “randomForest” package. The algorithm is based on conditional inference trees and applies subsampling without replacement, which was shown to produce reliable variable importance measures when predictors varied in their scale of measurement or number of categories.

Random forest regression models were fit using each site-year as a training set. The hybrid/variety variable was excluded as a predictor because predictions could not be made across site-years for which different hybrids/varieties were sown. The models may therefore be described as “agnostic” to the hybrid/variety planted. Fitted models were then used to predict yield for each remaining site-year planted with the same crop as the test set. If the soil type for a given experimental unit grid cell in the test set was not observed in the training set, soil type for that grid cell was set to missing. Prediction accuracy was assessed as the Pearson’s correlation between the predicted and observed values for yield. Once prediction accuracy had been assessed for all pairs of site-years, the mean prediction accuracy was calculated for each site-year as the training set using its accuracies for all other sites-years as the test sets.

Variable importance measures for each predictor were calculated for each site-year using the “varimp” function in the “party” package for R. Briefly, variable importance is calculated by permuting each predictor variable to determine the difference in prediction accuracy before and after the permutation. Variable importance measures were scaled to enable comparisons across site-years. To serve as a metric for the similarity between pairs of site-years, Euclidean distances between site-years in terms of their scaled random forest variable importance measures were calculated using the “dist” function in R.

A subset of the full dataset was utilized to evaluate the effect of the *mtry* hyperparameter on model accuracy (data not shown). In summary, prediction models were trained for each site-year using all possible values for *mtry* {1…*p*}. The fitted models were then applied to the remaining site-years in the subset planted with the same crop. Overall, prediction accuracies showed minimal variation among the evaluated *mtry* values. For each fitted model, there was no *mtry* value that consistently provided the highest accuracies across the predicted site-years. Therefore, the default *mtry* value of *p*/3 was used for this study (Breiman, 2001). The *ntree* value was set to 1000.

### Optimized Planting Rate Designs

Optimal planting rate designs were developed for each site-year using its own data as the training set to fit the random forest regression model. For each experimental unit grid cell, trained models were used to predict yield at each of the four tested planting rates, given the underlying soil and topographical features of that grid cell. The planting rate provided the highest yield prediction within each grid cell was then identified to create the optimized design.

## RESULTS

### Grain Yield Summary Statistics

The average grain yields across all site-years were 11,566 kg ha^-1^ for corn and 3,676 kg ha^-1^ for soybean, which were above the average statewide yields of 9,953 and 3,026 kg ha^-1^ for corn and soybean, respectively, (USDA-NASS, 2019) during the 2014-2018 period. Corn yields were lowest and most variable during the 2015 growing season, which was characterized by extremely wet conditions during May and June followed by dry conditions from early July through mid-September (Fig. 2). Scouting reports recorded in early July noted multiple sites with standing water in some low-lying sections (data not shown). Corn yields were likewise variable in 2016, due to severe drought conditions in June and July in western and central New York (Fig. 2). For soybean, 2015 yields were the most variable with respect to within-site-year standard deviation, while yields were the lowest in 2016.

### Effect of Planting Rate and Hybrid/Variety on Yield

Although ANOVA of planting rate showed a significant difference in yield between at least one pair of planting rate levels (*p*-value < 0.05) for 31 out of 32 site-years for corn, planting rate on average explained only 4.1 percent of yield variation and did not exceed 17.4 percent for any site-year for corn (Fig. 6). For soybean, a significant (*p*-value < 0.05) treatment effect for planting rate was observed in all 25 site-years, but the average amount of yield variation explained was likewise low at 10.8 percent. In contrast with the corn trials, however, planting rate explained a large amount of yield variation in some site-years (up to 40.8 percent).

**FIGURE 6.**
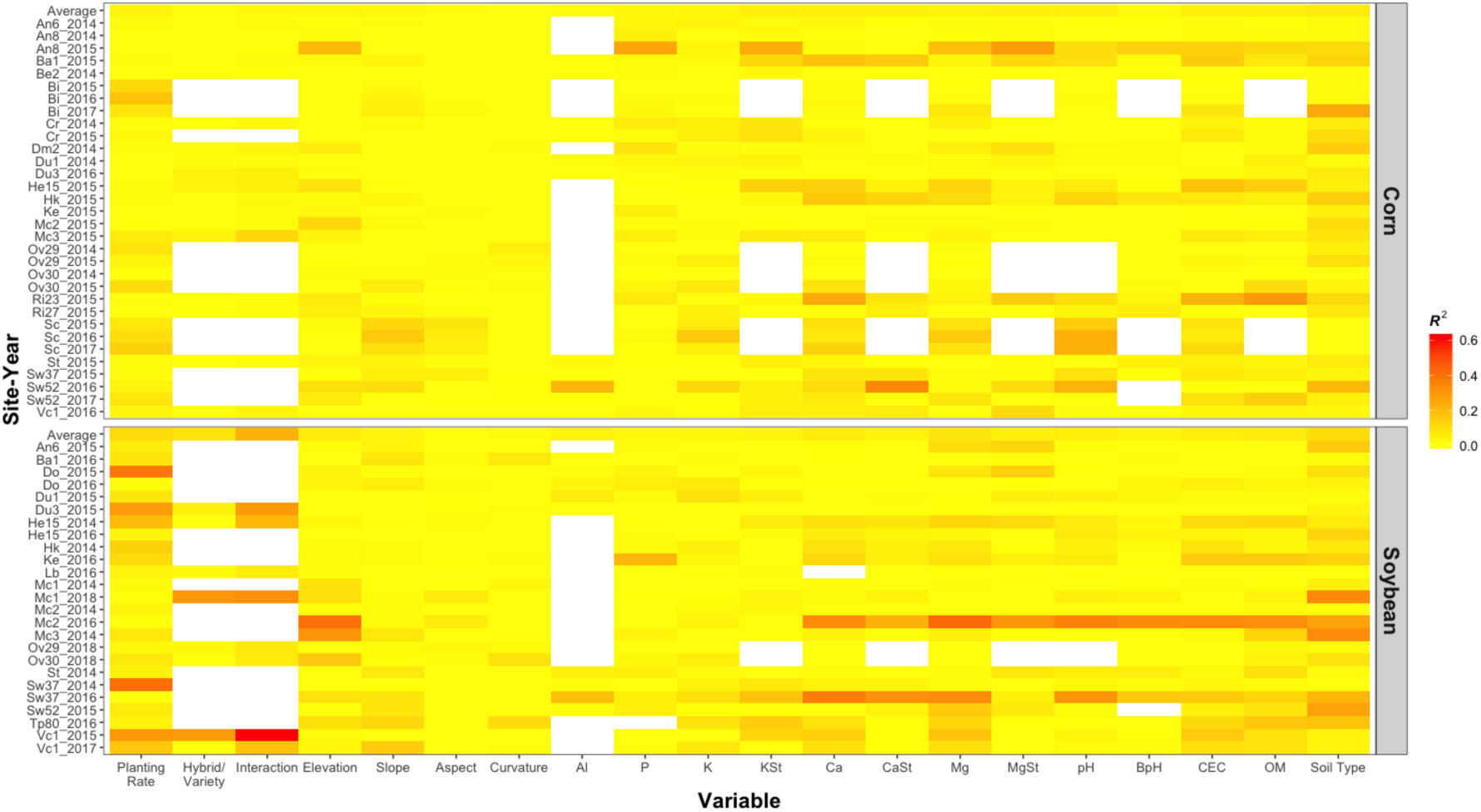
*R^2^* values of yield regressed onto each variable in each site-year with simple linear regression “Interaction” refers to the *R^2^* of yield regressed onto planting rate, hybrid/variety, and their interaction. The top row for each crop is the average over all site-years of that crop. White indicates that the variable was not measured or could not be included in the analysis due to low spatial autocorrelation.

Significant treatment effects (*p*-value < 0.05) were less prevalent for hybrid/variety, appearing in 13 out of the 19 site-years for which multiple hybrids of corn were sown and for 5 out of 9 site-years for which multiple varieties of soybean were sown. As with planting rate, the amount of yield variation explained by the hybrid variable was very low for corn, averaging 1.0 percent with a range of 0.0 to 4.1 percent. For soybean, variety explained 8.5 percent of yield variation on average, though this ranged as high as 30.1 percent. Notably, hybrid/variety explained over 25 percent of variation in soybean yields in 2 site-years, Vc1_2015 and Mc1_2018, in which the difference in performance between the varieties grown were 771 and 395 kg ha^-1^, respectively.

Considering the composite effects of planting rate, hybrid/variety, and their interaction, as before, the percent of yield variation explained was much greater for soybean than corn. These variables explained 2.8 and 22.8 percent of yield variation in corn and soybean, respectively. Most notably, these variables explained 61.9 percent of the variation in yield for Vc1_2015 with both varieties P22T41 and P92Y51 showing positive linear responses to planting rate.

### Topographical and Soil Summary Statistics

The 27 field sites tested varied greatly in their topographical and soil feature profiles. The lowest and highest elevations recorded were 105 and 368 masl, and the difference between lowest and highest points at a site was 16 m on average. The largest gradient within a single site with respect to elevation was observed at Ke, which ranged from 116 to 164 masl for a difference of 48 m. The United States Environmental Protection Agency classifies crop production on slopes ≥ 9 percent, or 5.14°, as “agricultural land cover on steep slopes.” Of the 27 field sites, 19 contained areas of sloped terrain at grades of 5.14° or above.

Soils in New York State are generally acidic. The soil grid sampling data showed that the pH ranged from 5.8 to 7.2 for the tested field sites. High levels of aluminum were likewise observed, ranging from 657 to 767 ppm. CEC levels were moderate, ranging from 6.2 to 16.5 meq/100 g, while soil OM was moderate to low, averaging 2.3 percent. According to the NRCS Soil Survey, 48 unique soil types were present across the tested sites, with 13 appearing in only one site. Each site contained an average of 6 different soil types, with type “Cazenovia” as the most common, appearing in 10 of the sites.

In the context of yield prediction, it would be desirable for the correlation for a pair of topographical or soil characteristics to have a high magnitude mean and low standard deviation across the testing sites, indicating that the two variables are not only strongly related, but their relationship is consistent across sites. A wide variety of relationships were observed between the measured variables (Fig. 7). As expected, pH and Al were observed to have a consistent, strong negative correlation, while CEC and OM were positively correlated. The relationship between OM and elevation was observed to be slightly negative, on average, although highly variable across the sites. When considering each variable’s relationship with yield, soil type had the most consistent relationship for both corn and soybean. It explained an average of 6.8 and 10.4 percent of the variation in corn and soybean yields, respectively, across the site-years. All other variables typically explained moderate amounts of yield variation within some site-years, but these relationships were not consistent across site-years overall.

**FIGURE 7.**
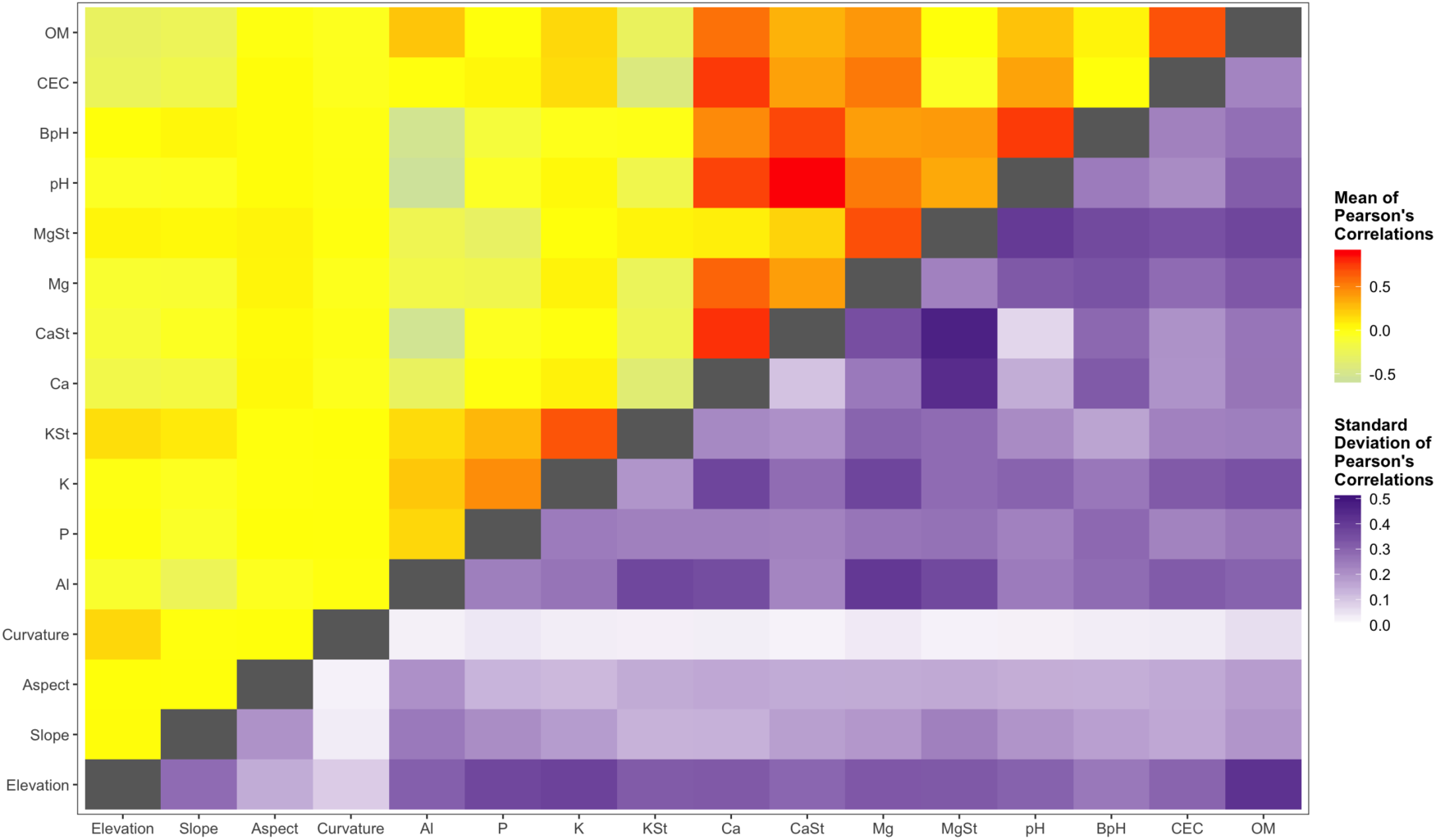
Means and standard deviations of the Pearson’s correlations between each pair of variables across the 27 field sites

### Random Forest Regressions

Random forest regression models were fit to predict yield using each site-year as the training set with moderate success. Overall, within site-year prediction accuracy was moderate, with *R^2^* values ranged from 0.16 to 0.78 and averaging 0.47 for corn and ranging from 0.17 to 0.72 and averaging 0.50 for soybean. Planting rate consistently had higher values for random forest regression variable importance for both crops but particularly for soybean (Fig. 8). Elevation was likewise important for both crops in most site-years. Organic matter appeared to be more consistently important for soybean than for corn. The soil nutrient variables were more frequently of greater importance for corn than for soybean. For most other variables, the level of importance was inconsistent across site-years.

**FIGURE 8.**
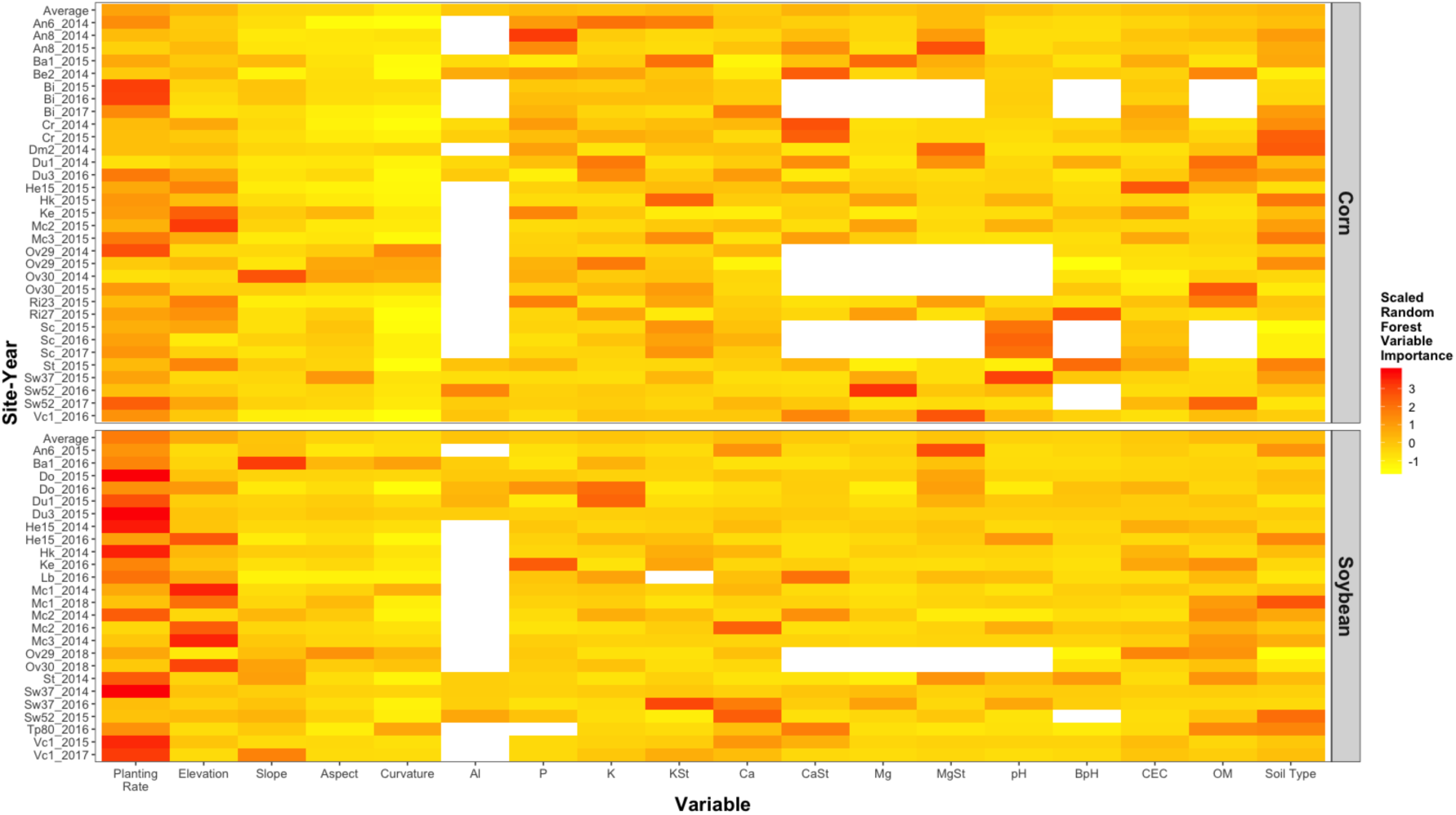
Scaled random forest regression variable importance measures for each variable in each site-year The top row for each crop is the average variable importance over all site-years of that crop, White indicates that the variable was not measured or could not be included in the analysis due to low spatial autocorrelation.

### Yield Predictions Across Environments

When random forest regression was applied to predict yield across site-years, the prediction accuracies varied greatly based on the pair of site-years used to train and test (Fig. 9). This is evident in the mean prediction accuracy for each site-year, which averaged 0.03 for corn and 0.08 for soybean. However, when considering pairs separately, each site-year predicted at least one other site-year at a level of ≥ 0.20 (with the exception of site-year Ba1_2016). The site-years with the highest mean prediction accuracies across all other site-years were Mc3_2015 for corn at a level of 0.11 and Du3_2015 for soybean at a level of 0.20. The site-year pairs with the highest accuracies were generally those for the same field site tested during different years. For example, site-year Bi_2015 had an average prediction accuracy of 0.05 across all site-years but predicted Bi_2016 with an accuracy of 0.64. For field sites that were tested for the same crop under two different years, the target planting rate design was re-randomized from one year to the next with the exception of Sc_2015 and Sc_2016. Those two site-years were sown using the same planting rate randomization in both 2015 and 2016. Notably, Sc_2015 and Sc_2016 predicted each other with accuracies of 0.76 and 0.75, respectively. Slightly lower accuracies of 0.63 and 0.65 were observed, respectively, when predicting Sc_2017, which was planted under a re-randomized design. The prediction accuracies for each pair of site-years were correlated with the Euclidian distances between the variable importance measures for each pair of site-years at a level of −0.23 for corn and −0.48 for soybean, suggesting that site-year pairs with similar variable importance profiles tend to predict one another at a higher level of accuracy.

**FIGURE 9.**
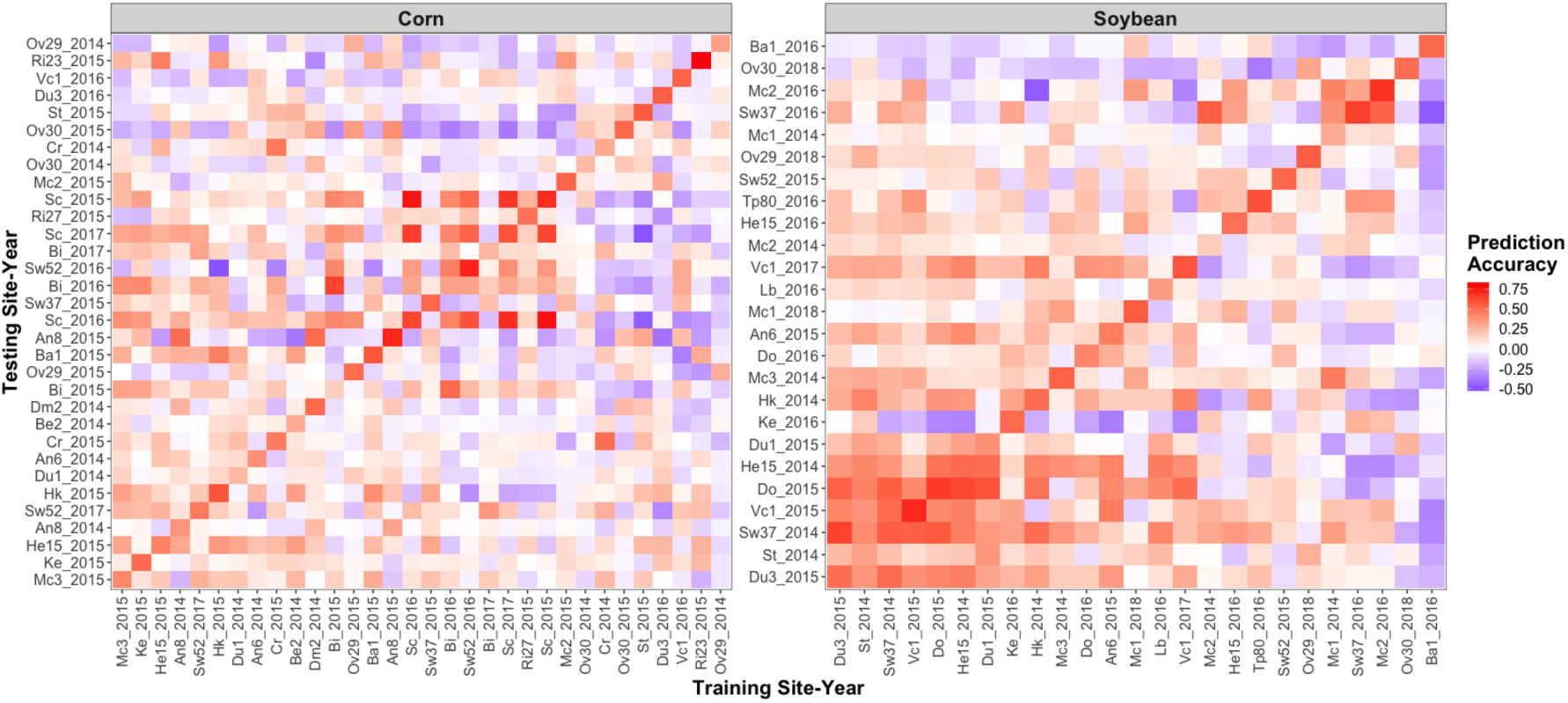
Across site-year prediction accuracy of random forest regression models Site-years are ordered with respect to their mean prediction accuracy across all other site-years. The diagonal corresponds to the *R^2^* of the random forest regression training model for each site-year.

### Planting Rate Optimization

Optimized planting rate designs were developed from the random forest regression prediction models to maximize yields (Fig. 10). In general, the optimized designs favored higher rates for both corn and soybean. For corn, the four planting rates in increasing order were assigned to, on average, 18.6, 21.8, 33.3, and 28.2 percent of the field. For soybean, optimizing for yield overwhelming favored the highest rate, which was assigned to 60.1 percent of the field, on average, while the remaining three rates in increasing order were assigned to 9.7, 13.8, and 18.9 percent of the field, on average.

**FIGURE 10.**
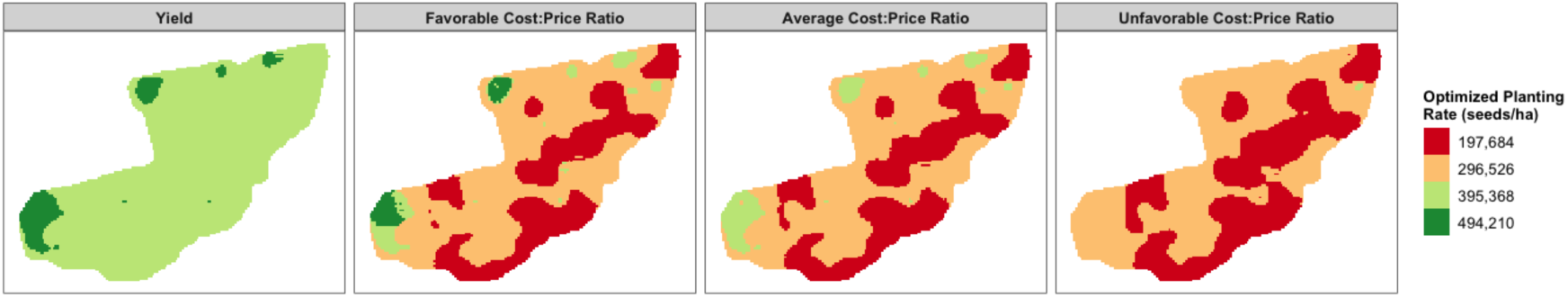
Example of the planting rate designs optimized yield and for profit at “favorable”, “average”, and “unfavorable” seed cost-to-market price ratios

Corn and soybean were evaluated in more than one year in 7 and 6 of the field sites, respectively. The optimized designs showed a considerable amount of differences depending on which year was used as the training set. For corn, the proportion of grid cells assigned to the same rate across years was 31.5 percent on average, ranging from 12.7 to 44.3 percent. For soybean, the average was 33.2 percent with a range of 5.3 to 74.3 percent.

## DISCUSSION

In the context of linear modeling, planting rate explained relatively low levels of the yield variation. In most site-years, there was no consistent yield advantage of one of the tested planting rates over the others. This lack of an apparent site-wide “optimum” planting rate would suggest that subsections of the field may have different optimums, according to underlying field conditions. Furthermore, for random forest regression, which is capable of modeling complex and nonlinear interactions, planting rate ranked highly in terms of variable importance for both corn and soybean. Together, these results provide support for the hypothesis that there may be important complex interactions of planting rate with topographical and soil variables that are driving response in yield.

These observations also highlight the importance of selecting an appropriate modeling approach for large agronomic datasets. A similar study found artificial neural networks, which – like random forest regression – are capable of modeling nonlinear interactions, were superior to linear modeling approaches when predicting corn yields using soil and landscape features (Miao et al., 2006). Random forest regression is among a suite of established and emerging techniques in machine learning that are becoming more widely used in agricultural research to model complex systems (Henderson et al., 2005; Häring et al., 2012; Xiong et al., 2014; Chlingaryan et al., 2018; Liakos et al., 2018). As the amount and variety of environmental, climate, management, and economic data available to growers increase through innovations in precision agriculture, it will be important for the community to consider the limitations of linear modeling and to begin to leverage the advances made in machine learning-based approaches.

The proposed random forest regression model was extended to build optimized variable rate planting designs for maximizing yields. Marked differences were observed between corn and soybean with respect to the resulting optimized designs. For corn, although higher rates were generally favored, all four tested planting rates were assigned to considerable parts of the field in most of the optimized designs. For soybean, many of the optimized designs were predominantly assigned the highest of the four rates. While this may suggest that fixed rate planting may be more suitable for those sites, it is also important to consider the trade-off between marginal yield gains from higher planting rates and the cost of the seed, as it would be no longer profitable to plant at higher rates if revenue from the yield gains sold at the market price does not exceed the cost of the additional seed. The proposed random forest-based approach may also be utilized to optimize planting rate designs for maximized profit. An example of optimized designs based on theoretical seed cost-to-market price ratios is shown in Fig. 10, demonstrating that lower planting rates are assigned to greater proportions of the field as the cost-price ratio becomes less favorable to growers. This highlights the importance of leveraging market trends to inform management decisions concerning variable rate, and the proposed approach is fully capable of accommodating user-defined commodity price and seed cost information to optimize planting rate designs for profit.

For sites that were evaluated under the same crop in multiple years, the resulting optimized planting rate designs showed considerable differences depending on the data used to train the model. These inconsistencies are likely due to variations in environmental conditions across years (Wells, 1993; Reeves & Cox, 2013) or differential sensitivities to planting rate among the varieties sown (Agudamu et al., 2016). Year-to-year variation in weather conditions and its impact on yield pose a challenge for the development of variable rate planting strategies. With sufficient training data, weather information could be incorporated into variable rate planting models, as with many of the currently available variable rate input management schemes (Melkonian et al., 2008, Hedley & Yule, 2009). However, this may be of limited use because, whereas variable rate input management strategies can be designed to respond to in-season weather status, planting decisions must be made at the beginning of the season without prior information about eventual weather outcomes. Therefore, an alternative approach may be to train models with data characterizing the range of possible weather scenarios in order to identify and base variable rate planting designs on main effects. Further long-term testing is needed to sample a greater range of seasonal weather conditions and to assess the stability of variable rate planting optimizations across years.

Short product life cycles also complicate the design and implementation of variable rate planting technologies. Results from this study and other works show that corn hybrids and soybean varieties can differ in their response to planting rate (Rutger and Crowder, 1967; Modarres et al., 1998; Agudamu et al., 2016). To address this, some seed companies conduct planting rate trials across a range of environments and publish optimal rate recommendations for the hybrids and varieties they release. This information cannot, however, be customized to the conditions of an individual site, nor does it provide insight into effective variable rate strategies to impose within a site. Empirical testing of variable rate response, as in this study, is time-consuming, and advances in agricultural biotechnologies and molecular breeding have accelerated the rate at which improved hybrids and varieties enter the market. A 2010 study estimated that US corn hybrids reach maximum sales with 2-3 years before declining in the following years (Magnier et al., 2010). New hybrids or varieties may therefore become obsolete during the time required to conduct thorough on-farm trials of variable rate response if carried out by the grower. Greater integration of variable rate testing in the germplasm development pipeline may serve to address this issue. Alternatively, though not ideal, variable rate optimization models could be trained using data collected on a range of hybrids/varieties so as to become “agnostic” the hybrid or variety sown and base planting rate optimizations on main effects. Empirical evaluations are needed to establish the value of variable rate planting if hybrid/variety effects are ignored.

In addition, the expense of conducting on-farm trials of variable rate response is not trivial. Grid soil sampling was carried out at each field site, and research associates trained in data science were needed to process and analyze the data. In addition, the current approach requires growers to plant and evaluate variable rate randomized block designs for at least one year before optimized designs can be generated. The ability to build optimized designs for sites that have not undergone variable rate randomized block testing would therefore be of great value. Overall, using the fitted random forest regression models to predict yield across site-years produced low prediction accuracies. The site-years with the highest mean prediction accuracies across all other site-years for corn and soybean were Mc3_2015 and Du3_2015 with accuracies of 0.11 and 0.20, respectively. Low accuracies are perhaps expected given the variation across site-years in the direction and magnitude of the relationships between predictor variables. Within site-year prediction accuracies were much higher, indicating that local randomized block testing is likely to provide the most accurate optimizations; however, there may be alternative approaches for building the most useful training sets for a given untested site. For example, there was a significant negative correlation between the across site-year yield prediction accuracies and the Euclidean distances between site-years for random forest regression variable importance. Provided yield, topographical, and soil data are available for an untested site, random forest regression models predicting yield could be fit, and the resulting variable importance measures could be utilized to identify the most similar site-year(s) evaluated under the randomized block design to use as a training set. Additionally, data from multiple site-years could be combined in an optimal fashion to maximize prediction accuracy. Heslot et al. (2013) provides an innovative approach for this type of training set optimization in the context of plant breeding that identifies and removes less predictive site-years from the complete set of site-years in order to train a composite model.

Prior to testing planting rate optimization for untested sites, a critical next step is to empirically validate optimized designs based on local testing. Within site-year prediction accuracies suggest that the data collected are capturing the relationships between yield, planting rate, topography, and soil at a reasonable level, but in order to determine whether the proposed approach offers advantage over fixed rate sowing across years, empirical testing of the optimized designs is needed. The NYCSGA is currently conducting on-farm validation trials in which the optimized designs developed from previous years’ randomized block variable rate trials are sown.

## CONCLUSIONS

In this study, we have proposed a random forest regression-based approach to predict yields given high resolution topographical, soil, and variable rate planting rate data. We have likewise developed a method for optimizing planting rate designs to maximize yields, thought it may be further extended to accommodate user-defined seed costs and crop market prices for maximizing profits. While planting rate explained relatively low amounts of yield variation in the linear context, it ranked highly in terms of random forest regression variable importance, suggesting that there may be important complex, non-linear interactions of planting rate with topographical and soil features driving yield response that are not captured by linear model. The resulting optimized planting rate designs showed limited consistency across years, likely due to temporal weather variation as well as changes in the hybrid/variety sown. Further long-term studies will be necessary to evaluate the stability of optimized designs across years and for varying hybrids/varieties. Lastly, while the ability to predict yield within site-years was moderate, predictions accuracies across site-years were low, perhaps due to the wide variability of the magnitude and direction of the relationships between topographical variables, soil features, planting rate, and yield. Local testing will most likely provide the most accurate planting rate optimizations; however, further research is needed to evaluate strategies for the creation of training sets to build optimized variable rate planting designs untested sites. On-farm validation experiments of the proposed approaches are ongoing.

## ACKNOWLEDGEMENTS

This project was designed and administrated by the New York Corn and Soybean Growers Association. Research funding was provided by the New York Corn and Soybean Growers Association and the New York Farm Viability Institute. Additionally, this work was supported by Hatch funding from the USDA National Institute of Food and Agriculture. We are also thankful to the New York Soybean Checkoff, DuPont Pioneer, and the National Science Foundation Graduate Research Fellowship (Grant No. DGE-1650441) for supporting the graduate studies of Margaret R. Krause. The authors express appreciation to all New York State corn and soybean growers who contributed their time, land, and resources for the on-farm field experiments.

---

**FIGURE S1** The pipeline for aggregating the target planting rate, as-applied planting rate, yield, soil survey, and grid soil sampling to a common grid and performing quality control

**A** Removing the headlands from the as-applied planting rate and yield data layers

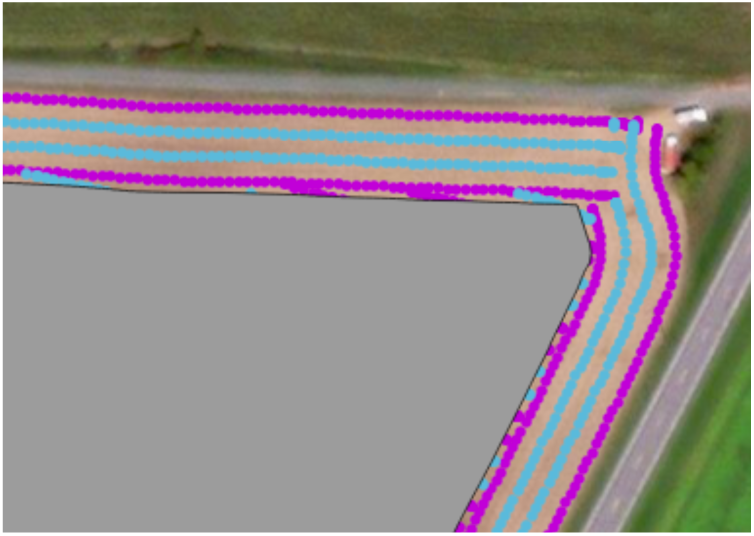

A buffer was created to exclude the headland areas using the “Buffer” function in QGIS.

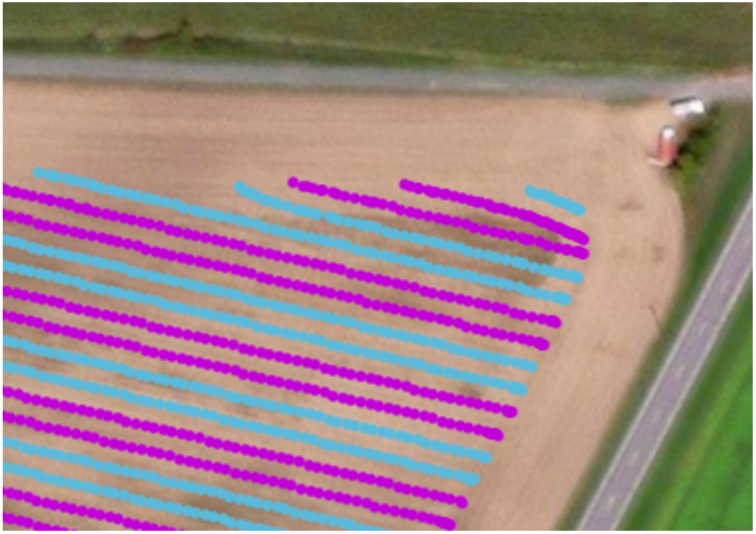

The headland areas were removed from the as-applied planting rate and yield layers using the “Clip” function in QGIS.

**B** Removing grid cells based on number of as-applied planting rate or yield data points per cell

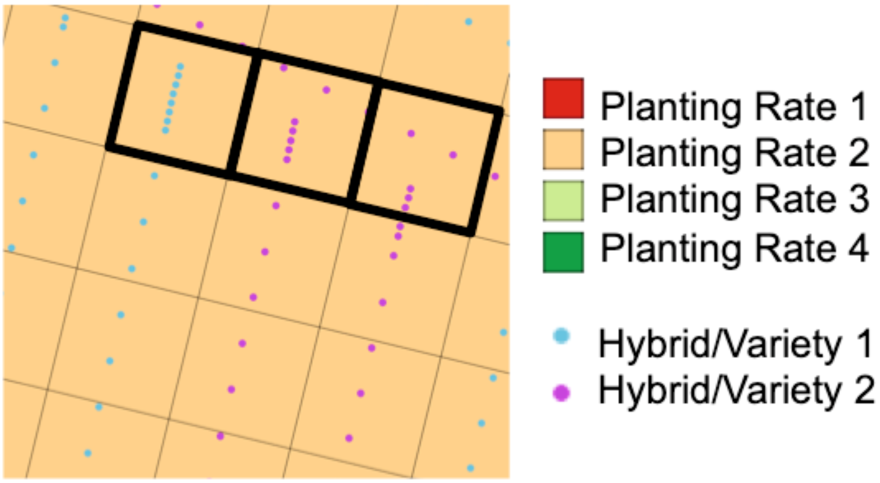

Along the edges of the field in particular, the planter and combine often record many data points in rapid succession. To reduce the influence of these data on the analysis, each experimental grid cell was assigned a unique ID. The as-applied planting rate and yield data points within each cell were assigned the respective experimental grid cell ID in QGIS. The number of data points within each grid cell was calculated in R. Significant outliers were identified using Studentized residuals (*p*-value < 0.05) with the lm() and rstudent() functions. Experimental grid cells identified as outliers with respect to the as-planted or yield data layers were removed from the analysis. In the illustrated example, the cells outlined in black boxes would have been identified as outliers and removed from the analysis.

**C** Removing grid cells that contain data from more than one target planting rate

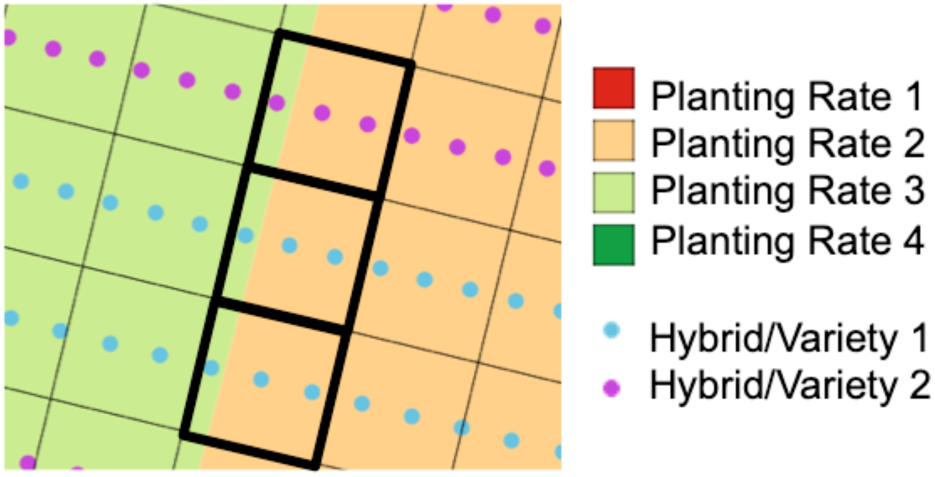

Because the 0.81 ha target planting rate blocks were not always proportional in size to the experimental unit grid cells, it was not possible to precisely align the experimental unit grid cells with the target planting rate blocks. This resulted in some grid cells containing data corresponding to more than one target planting rate. Using the unique IDs matching data points to grid cells, the target planting rate for each data point within each grid cell was identified in R. Cells that contained data points corresponding to more than one target planting rate were removed from the analysis. In the illustrated example, the cells outlined in black boxes would have been identified and removed from the analysis.

**D** Removing grid cells that contain data for more than one hybrid/variety

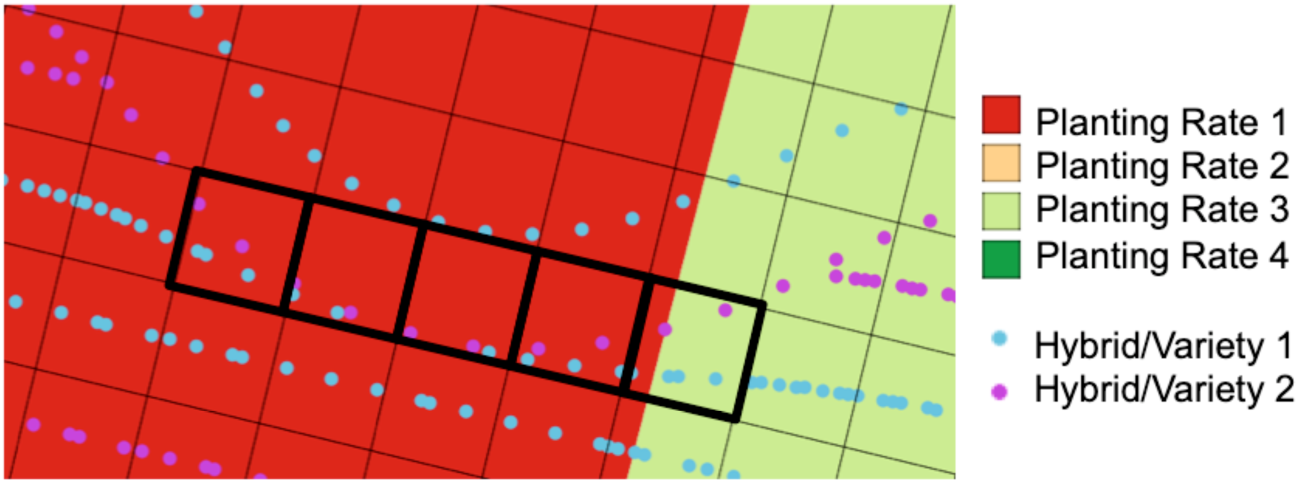

Due to the shape and features of each field, the planters occasionally deviated from a straight line path, resulting in data corresponding to more than on hybrid/variety falling within the same experimental unit grid cell. Using the unique IDs matching data points to grid cells, the hybrid/variety sown for each data point within each grid cell was identified in R. Cells that contained data points corresponding to more than one hybrid/variety were removed from the analysis. In the illustrated example, the cells outlined in black boxes would have been identified and removed from the analysis.

**E** Removing outlying grid cells based on the mean and variance of the as-applied planting rate and the yield

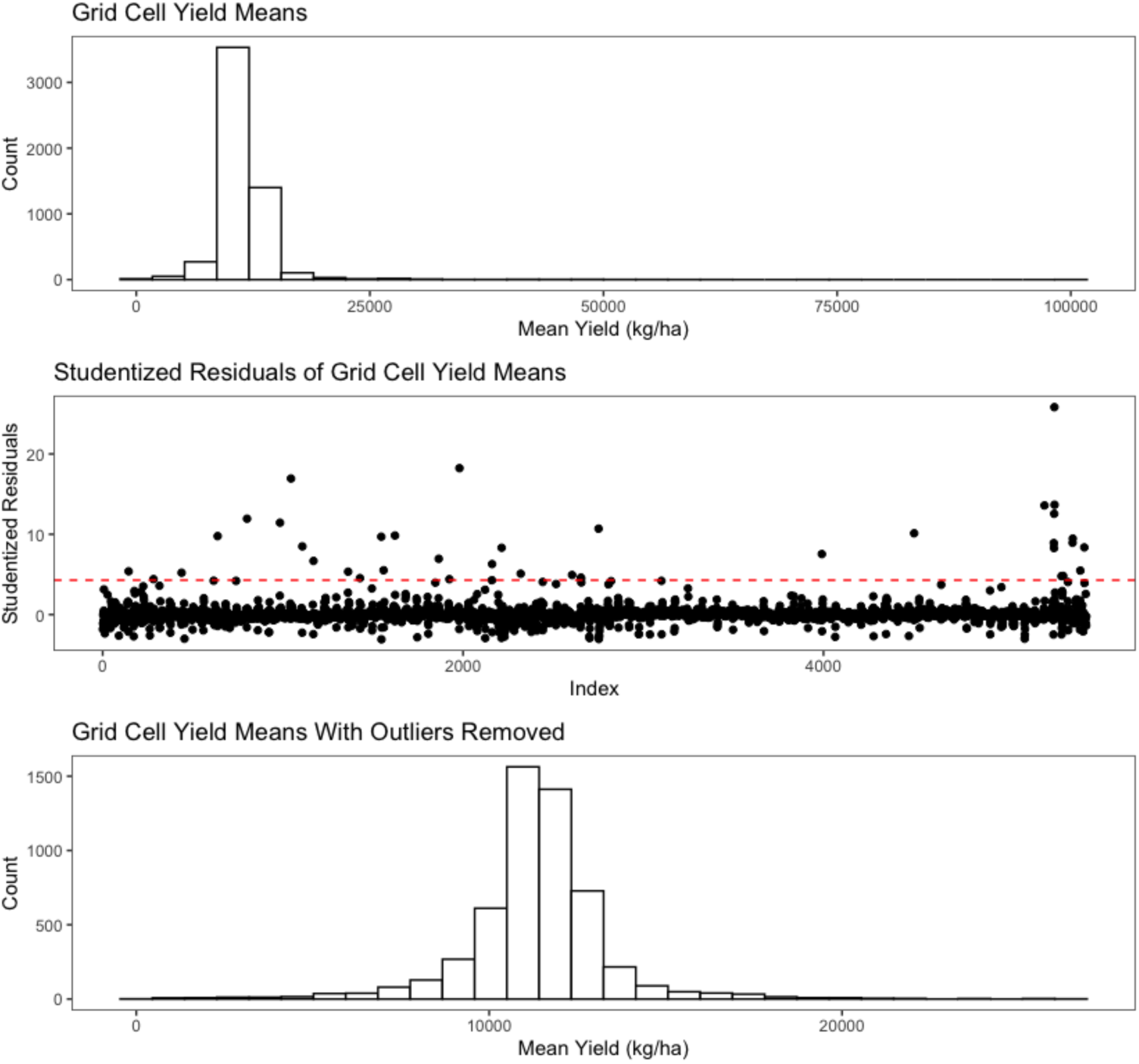

It was not uncommon for the planter and combine to record extreme values for the as-applied planting rate and the yield. To reduce the influence of these data points on the analysis, the mean and variance of the as-applied planting rate and yield for the data points within each grid cell were calculated in R. Significant outliers were identified using Studentized residuals (*p*-value < 0.05) with the lm() and rstudent() functions. Experimental grid cells were removed from the analysis if they were identified as outliers with respect to the mean or the variance of the as-applied planting rate or the yield. The illustrated example shows the distribution of yield means before and after outlier removal.

**F** Removing grid cells in which the as-applied planting rate did not correspond to the target rate

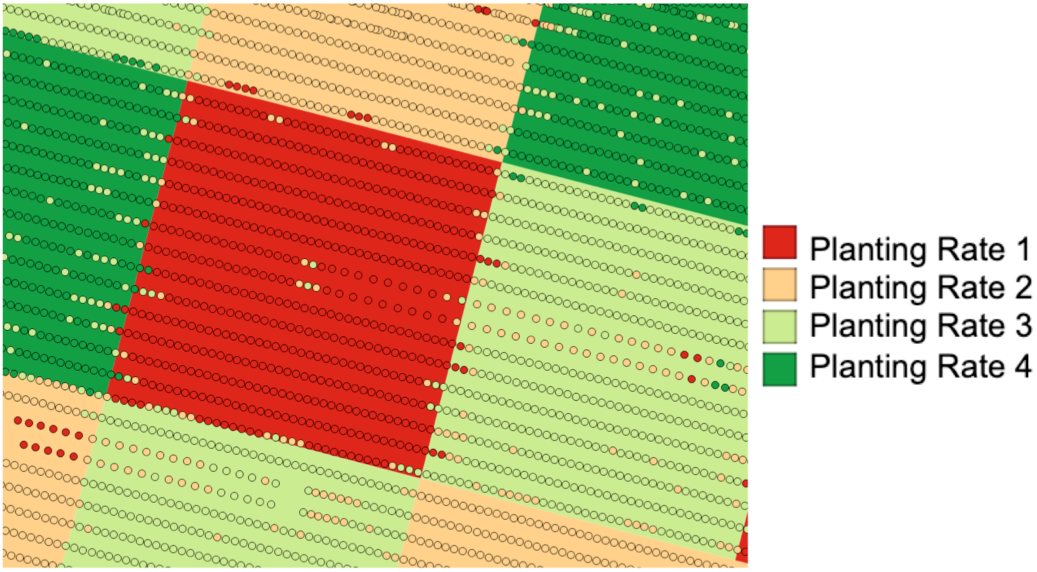

Occasionally, the as-applied planting rate did not match the target planting rate. In the illustrated example, the circles represent the as-applied data points and their colors indicate the as-applied rate. The blocks indicate the target planting rates.

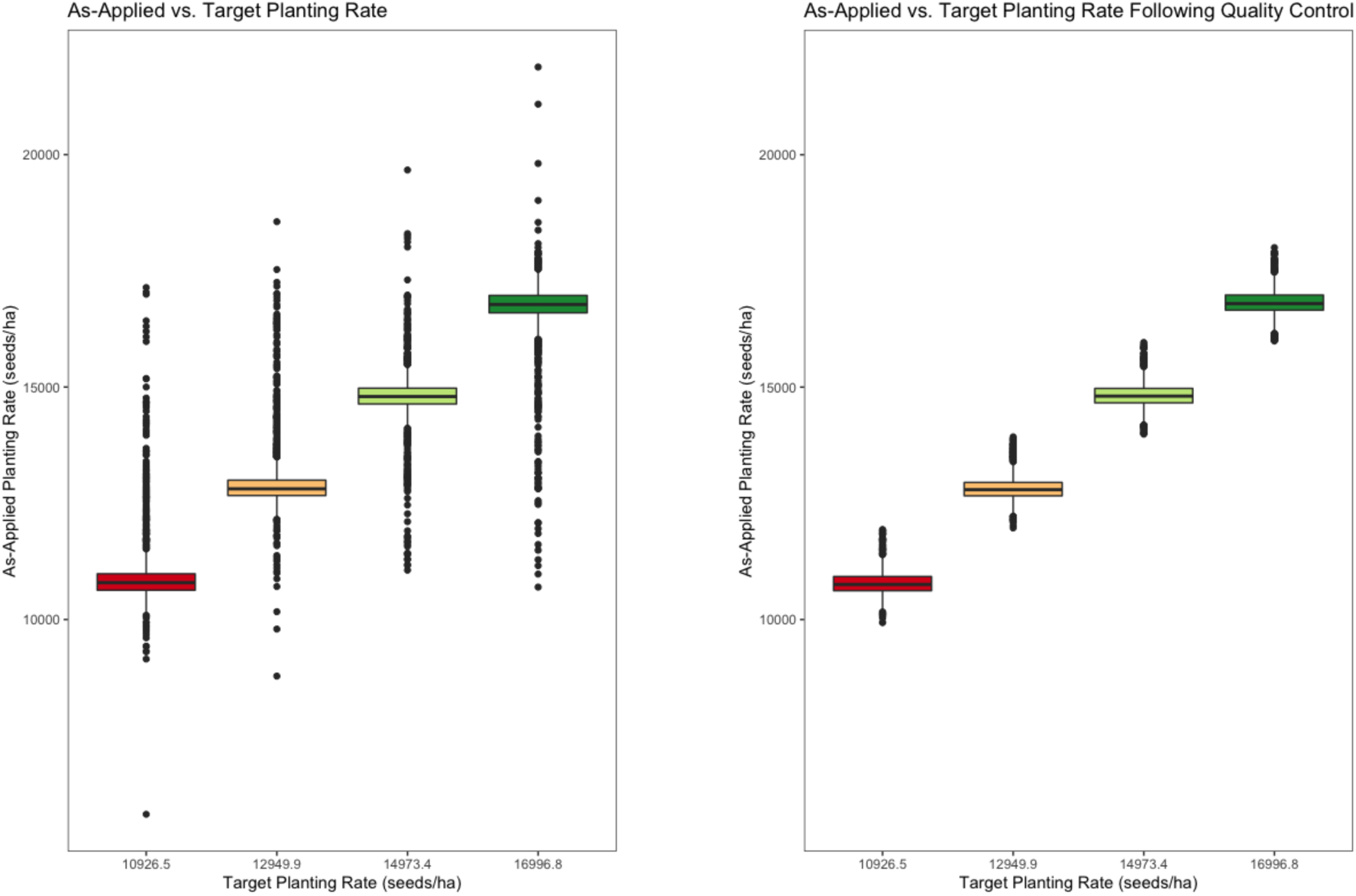

Experimental grid cells in which the as-applied planting rate data deviated substantially from the target planting rate were removed from the analysis. For corn, grid cells with as-applied planting rates that deviated from the target planting rate by > 1011.7 seeds ha^-1^ were removed from the analysis. For soybean, grid cells with as-applied planting rates that deviated from the target planting rate by > 8093.7 seeds ha^-1^ were removed from the analysis.

**G** Deriving the topographical variables from the as-applied planting rate layer

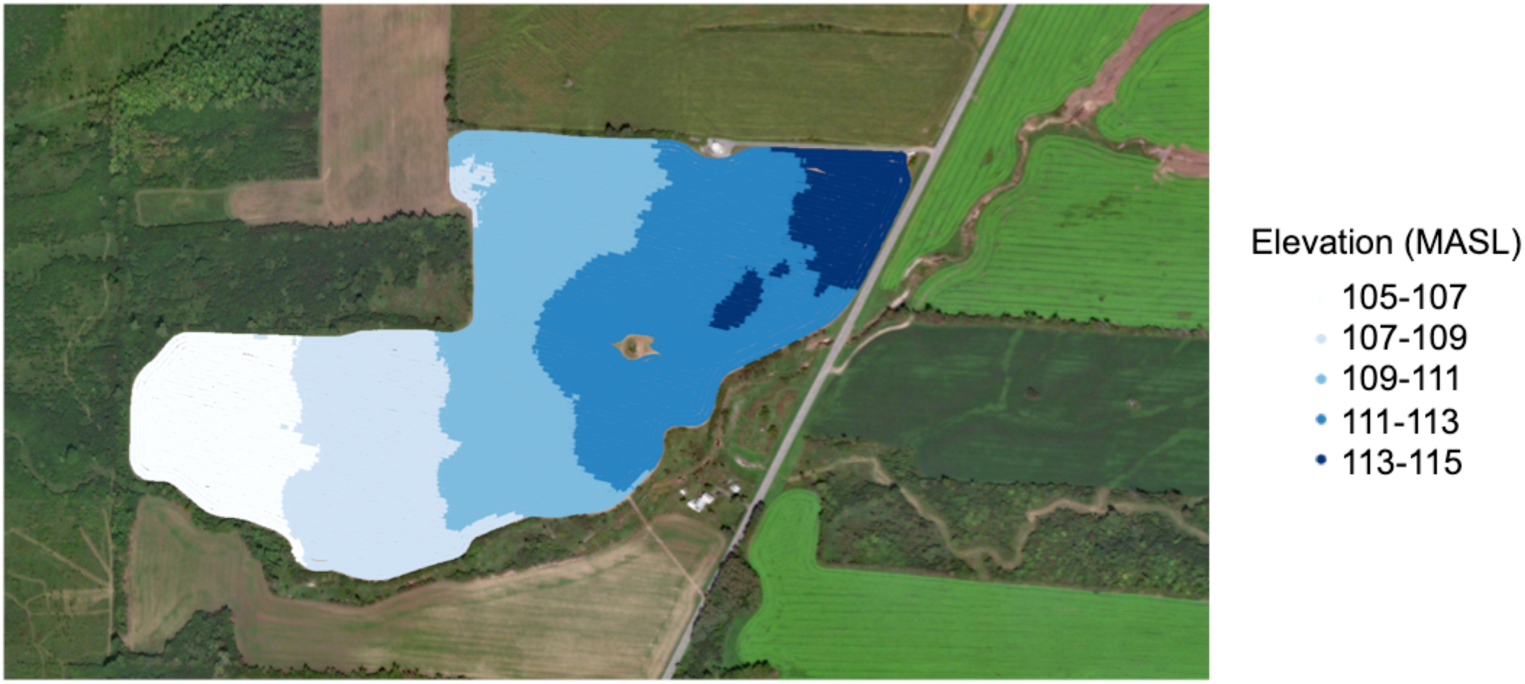

Elevation measurements were recorded with a GPS system aboard the planter.

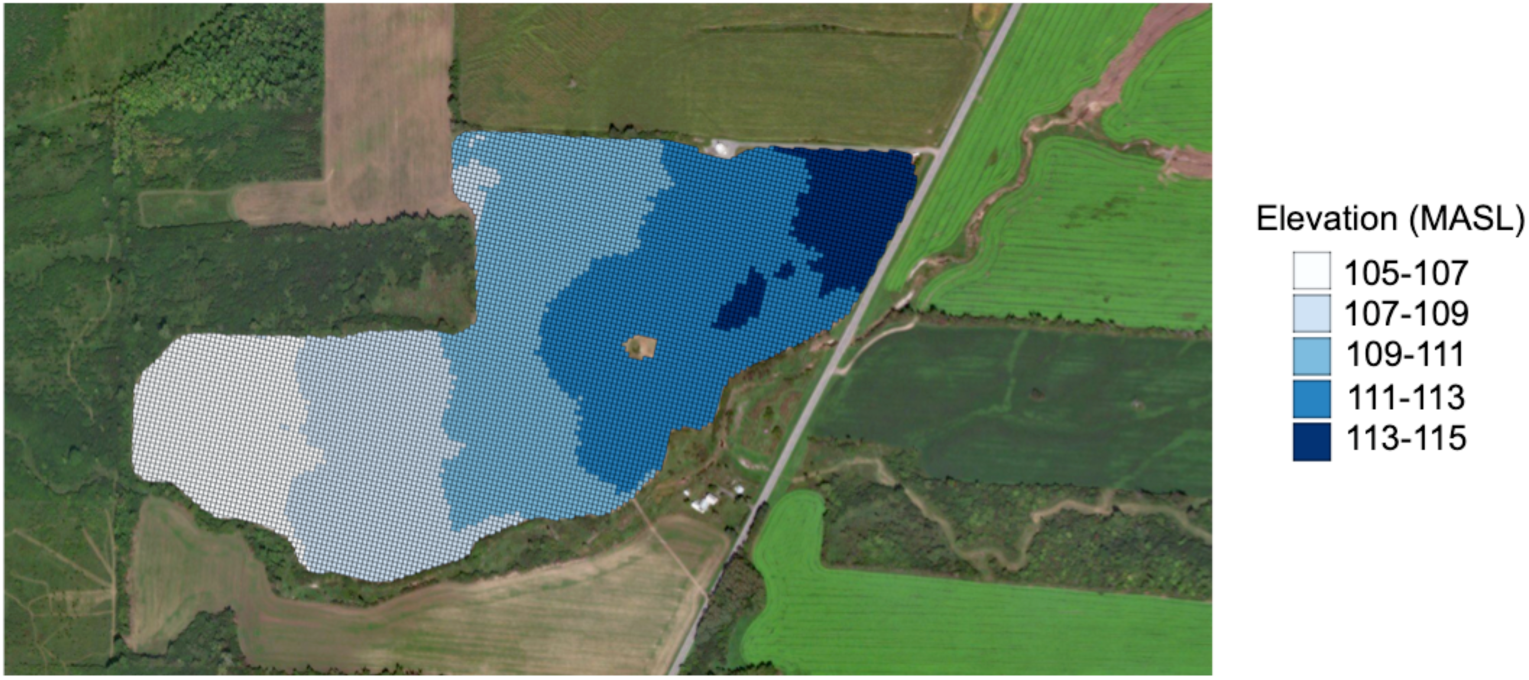

Means of the elevation data points within each experimental unit grid cell were calculated using the “Geoprocessor” function of the “HTP Geoprocessor” plugin in QGIS such that a single value for elevation is assigned to each grid cell.

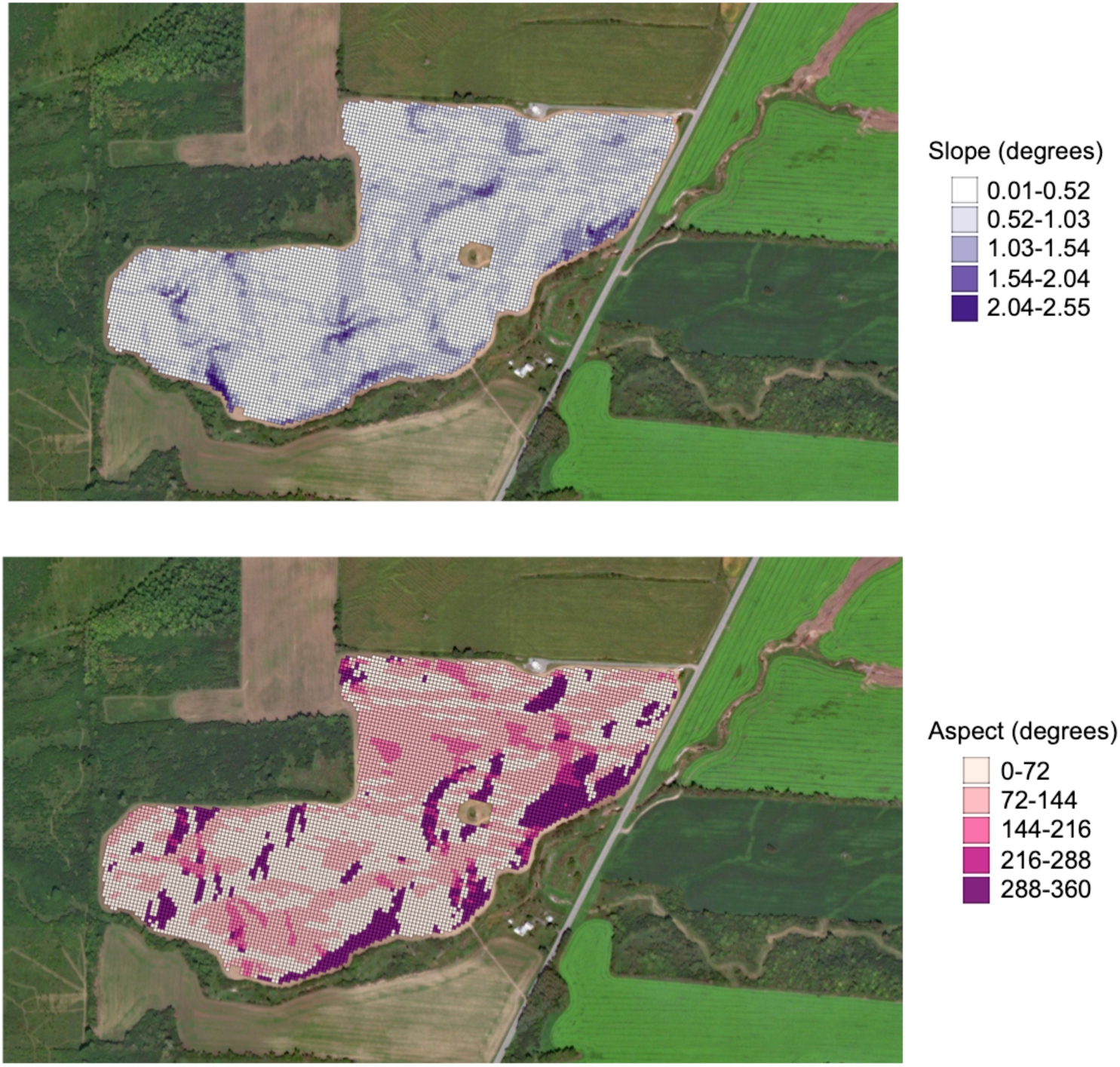

The slope and aspect for each experimental unit grid cell was calculated in R according to Burrough and McDonell (1998).

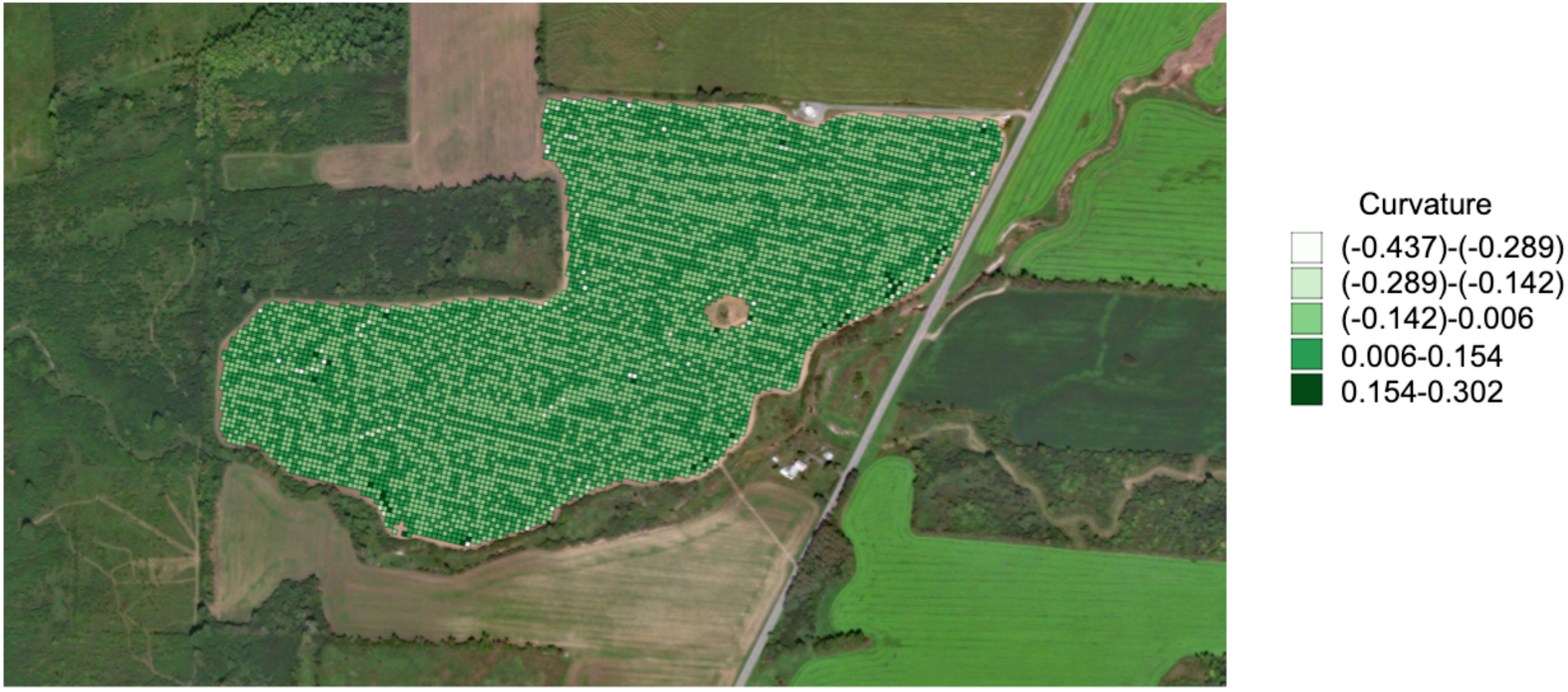

Curvature was calculated in R for each experimental unit grid cell according to Zeverbergen and Thorne (1987).

**H** Kriging the grid soil samples to the experimental unit grid

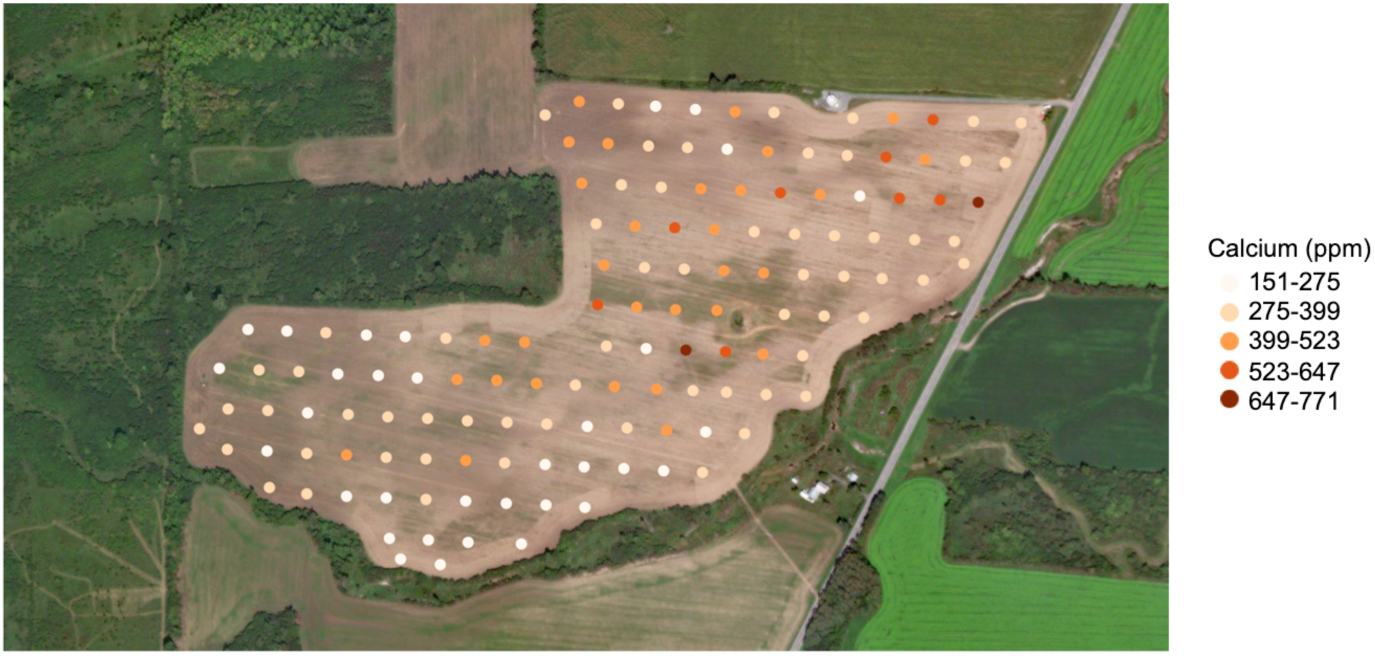

Soil grid samples were collected in a 0.20-ha grid pattern.

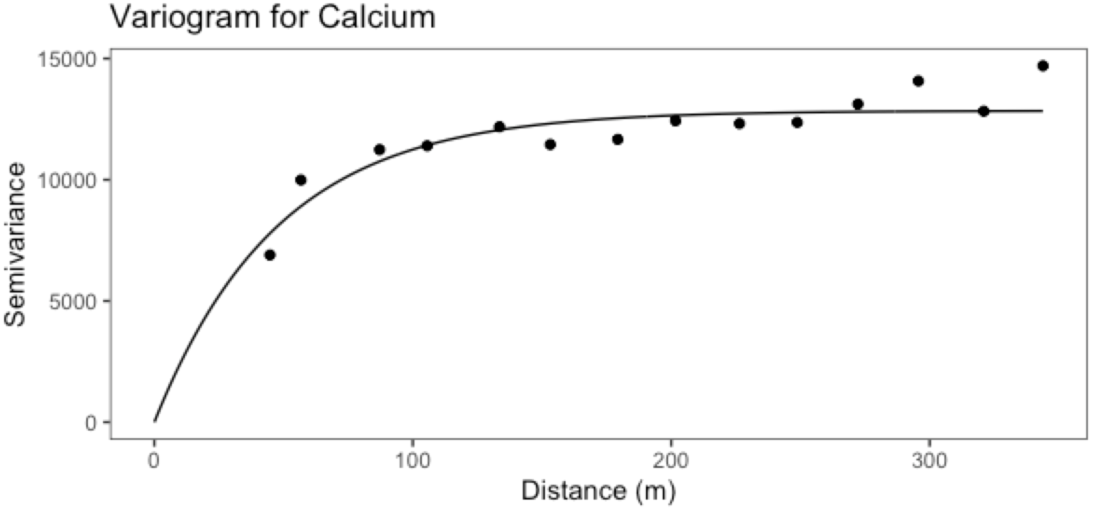

Using the “sp” and “gstat” packages in R, the variograms were fit using the fit.variogram() function for each of the 12 soil characteristics.

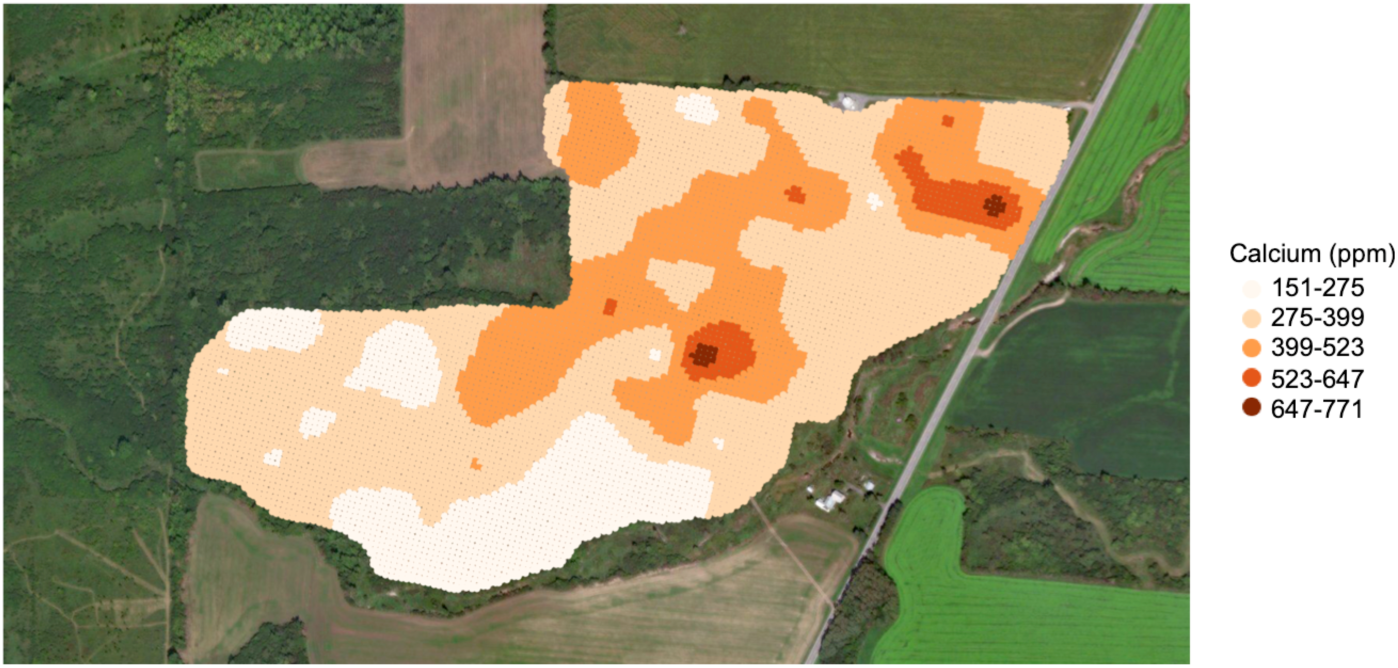

Centroids for each cell of the experimental unit grid were created using the “Polygon Centroids” function in QGIS. The krige() function from the “gstat” R packages was utilized to block krige the soil characteristics to grid. The “block” parameter set to the length and width of the grid cells.

**I** Setting the soil type to “missing” if more than one soil type was present within a grid cell

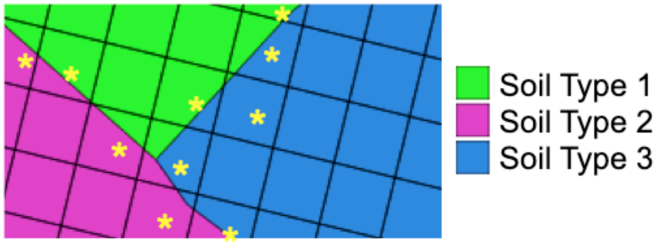

For experimental grid cells in which there was more than one soil type present, the soil type was set to missing. In the illustrated example, the yellow asterisks denote grid cells in which more than one soil type is present.

